# Learning to Control the Brain through Adaptive Closed-Loop Patterned Stimulation

**DOI:** 10.1101/2020.03.14.992198

**Authors:** Sina Tafazoli, Camden J. MacDowell, Zongda Che, Katherine C. Letai, Cynthia Steinhardt, Timothy J. Buschman

## Abstract

Stimulation of neural activity is an important scientific and clinical tool, causally testing hypotheses and treating neurodegenerative and neuropsychiatric diseases. However, current stimulation approaches cannot flexibly control the pattern of activity in populations of neurons. To address this, we developed an adaptive, closed-loop stimulation (ACLS) system that uses patterned, multi-site electrical stimulation to control the pattern of activity in a population of neurons. Importantly, ACLS is a learning system; it monitors the response to stimulation and iteratively updates the stimulation pattern to produce a specific neural response. *In silico* and *in vivo* experiments showed ACLS quickly learns to produce specific patterns of neural activity (∼15 minutes) and was robust to noise and drift in neural responses. In visual cortex of awake mice, ACLS learned electrical stimulation patterns that produced responses similar to the natural response evoked by visual stimuli. Similar to how repetition of a visual stimulus causes an adaptation in the neural response, the response to electrical stimulation was adapted when it was preceded by the associated visual stimulus. Altogether, our results show ACLS can learn, in real-time, to generate specific patterns of neural activity, providing a framework for using closed-loop learning to control neural activity.

## Introduction

The ability to control neural activity is an important tool for understanding the brain and for treating brain disorders. In neuroscience, causal manipulation of neural activity has been critical for testing hypotheses about how neural activity relates to behavior (e.g. decision making, Salzman et al., 1992; Tye et al., 2013) and for understanding how neural circuits function (e.g. the role of oscillations, Cardin et al., 2010). Clinically, stimulation of neural activity has emerged as a treatment option for neurological diseases: reducing tremors in Parkinson’s patients (Groiss et al., 2009), improving mood in patients with severe depression (Mayberg et al., 2005), and interrupting epileptiform activity (Zangiabadi et al., 2019).

However, current stimulation approaches are limited. Typically, only a few neurons are stimulated at time (e.g. with electrical stimulation, Histed et al., 2009) or populations of neurons are uniformly stimulated to enhance or suppress activity (e.g. with optogenetic stimulation; Cardin et al., 2010; Ritt and Ching, 2015). In contrast, neural representations are diverse and are distributed across the population. Therefore, interfacing with the brain for scientific or clinical applications requires a system that can independently control many stimulation sites to precisely modulate the activity of populations of neurons (Grosenick et al., 2015; Mardinly et al., 2018).

Precise, large-scale control of neural populations must overcome several obstacles. First, the complex, non-linear, and recurrent nature of the brain makes it difficult to predict the effect of stimulation on neural activity (Koch and Laurent, 1999). Previous work has suggested neural networks in the brain are semi-chaotic, such that small differences in inputs are amplified over time (Babloyantz and Salazar, 1985; Schiff et al., 1994). In addition, biological noise means the neural response to the same input can be variable (Faisal et al., 2008). Because of this, both optogenetic and electrical stimulation can have complex (often unintended) effects, both locally and in downstream brain regions (Allen et al., 2015; Wolff and Ölveczky, 2018).

Second, the brain is plastic. Functionally, this plasticity enables the brain to adapt to changes in the environment and one’s physiology. However, plasticity also changes how neurons respond to the same stimulation over time. This poses a challenge for chronic, clinical brain stimulation devices (Fallon et al., 2008). Similarly, changes in behavioral state (e.g. arousal) or the progression of disease can lead to global changes in the connectivity or firing properties of neurons, altering the response to stimulation over time.

Finally, individuals’ brains are unique. The localization of cognitive function varies significantly between individual mice, monkeys, and humans. This changes the effect of stimulation across subjects and can determine the efficacy of deep brain stimulation treatments (Greene et al., 2020). Individual differences in disease progression also mean the effect of stimulation will change over time in a unique way for each patient (Blauwendraat et al., 2017). The problem of individual differences is exacerbated as stimulation targets become more precise – the pattern needed to instantiate a specific cognitive variable in a neural population cannot be known *a priori*. Current stimulation approaches are unable to deal with this problem, as they rely on knowing the relationship between stimulation and response. For example, recent innovations enable recording and stimulation of large neural populations using all-optical recording/stimulation or multi-electrode recording/stimulation (Berger et al., 2012; Emiliani et al., 2015; Hampson et al., 2012; Marshel et al., 2019; Zhang et al., 2018). However, these approaches only work when one already knows what stimulation pattern will elicit a desired response: multi-electrode stimulation can compensate for a lesion, but it does so by reproducing the pattern of neural activity observed before the lesion occurred (Berger et al., 2011; Hampson et al., 2012). Yet, the relationship between stimulation and response is unknowable in many scientific applications and almost all clinical applications.

Altogether, the individualistic, non-linear and dynamic nature of the brain makes it difficult to rationally design stimulation patterns to control neural activity. This problem is not limited to neuroscience – control theory techniques work best when applied to simple, linear and nonlinear systems (Khalil, 2002). Indeed, control of neural systems has been successful when the structure of the system is well mapped (Ahmadian et al., 2011; Li et al., 2013), such as controlling the specific neurons responsible for locomotion in C. elegans (Yan et al., 2017). However, controlling complex systems, such as large-scale neural populations, is poorly understood (Ritt and Ching, 2015; Tang and Bassett, 2018).

One solution to this problem may be to use machine learning algorithms to learn to control nonlinear systems (Fleming and Purshouse, 2002; Rana and Zalzala, 1997). Here, we show that machine learning algorithms can learn to control the brain. To this end, we combined multi-site electrical stimulation with multi-electrode electrophysiological recording of neural activity. Then, we used a closed-loop machine learning algorithm to find the stimulation pattern that induced a specific pattern of activity in the neural population. Importantly, the algorithm does not need a model relating stimulation to response. Instead, it simply minimizes the error between the evoked response and its target response. We show the ACLS approach successfully learns to produce arbitrarily-defined target responses in a population of neurons. It is also robust to noise and drift in the response of neurons.

In this way, ACLS can overcome the obstacles in precisely controlling populations of neurons. First, by iteratively monitoring the effects of stimulation, ACLS does not need to predict how stimulation will affect the circuit. Second, because ACLS is constantly learning, it can compensate for dynamics that are on a longer timescale than its learning speed (e.g. slow drifts in state, plasticity and disease progression). Finally, the complexity and uniqueness of the brain is offset by the fact that ACLS can learn to control neural activity without needing to estimate a model relating stimulation to response.

Altogether, our results provide the first step in developing a system to causally manipulate populations of neurons. This will allow us to test open scientific questions relating neural population representations to cognition and lays the groundwork for improving clinical applications of brain stimulation.

## Results

Adaptive closed-loop stimulation (ACLS) works by iterating through three steps (Figure 1). First, the system observes the evoked response to a specific pattern of stimulation (Fig. 1A). Here, we measure the neural firing rates of a population of neurons. However, this response could be any measurable variable, such as distal neural activity, oscillations in the local field potential, BOLD signal, or behavior. Second, the ACLS evaluates an error function that determines how much the evoked response differs from the desired ‘target’ response (Fig. 1B). Here, we use Euclidean distance to a target pattern of neural activity, but ACLS could use different error functions to achieve other neural or behavioral states. Third, a machine learning algorithm updates the stimulation pattern in order to reduce the error value (Fig. 1C). By iteratively progressing through these steps, the algorithm learns a stimulation pattern that minimizes the error function and, thus, produces a desired response (see Box S1 for pseudocode).

**Figure 1.**
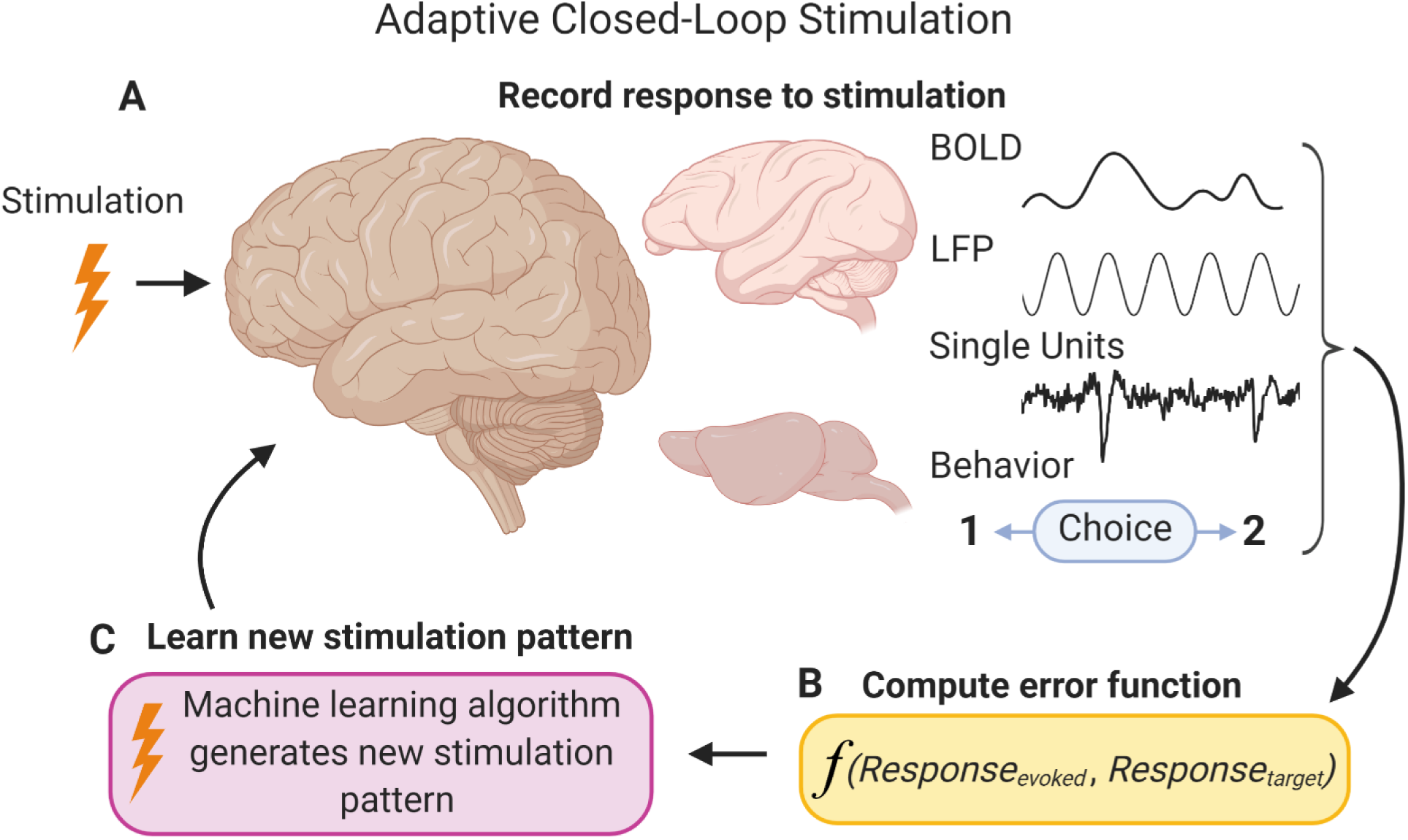
Schematic of Adaptive Closed-Loop Stimulation. Adaptive closed-loop stimulation (ACLS) works by iterating through three steps. **(A)** First, the system observes the evoked response to a specific pattern of stimulation. Here, we record multi-unit activity, but this could be any measurable variable (e.g. bold signal, local field potential, single units, or behavior). **(B)** Second, the ACLS evaluates an error function that determines the difference between the evoked response and the desired response. Here, we use Euclidean distance to a target pattern of neural activity, but this can be any error function (e.g. cosine similarity, correlation) to a target neural or behavioral state. **(C)** Third, a machine learning algorithm updates the stimulation pattern in order to reduce the error function. By iteratively progressing through these steps, the algorithm learns a stimulation pattern that minimizes the error function and, thus, produces a desired neural or behavioral response. ACLS pseudocode is supplied in Box S1.

### ACLS control of neural representations *in silico*

We first implemented ACLS *in silico*, testing its ability to control activity within a deep convolutional neural network (CNN). CNNs have been shown to approximate processing along the ventral visual stream and have representations that are similar to those found in the brain (Bashivan et al., 2019; Ponce et al., 2019; Walker et al., 2019; Yamins et al., 2014). Importantly, the CNN captures many of the properties of the brain (non-linearities and complexity) and so controlling it also faces many of the same obstacles as controlling the brain. This makes controlling CNNs a good starting point for testing ACLS.

We began by using ACLS to control a 5-layer CNN trained to classify 10 numeric digits from the MNIST database (Fig. 2, see Methods). ACLS delivered ‘stimulation’ inputs to the input layer of the CNN (i.e. a grayscale image, Fig. 2A). The ACLS learned the amplitude of stimulation input to each neuron in that layer (i.e. the pixel value, bounded in the grayscale range). The response of the CNN was taken as the pattern of activity in the last hidden layer (i.e. immediately preceding output; the ‘response’ layer in Fig. 2A). Similar to the brain, stimulus representations become more complex along the hierarchy of the CNN (Mahendran and Vedaldi, 2016). Indeed, the last layer of the CNN represented the categories of stimuli in a distributed manner, across the neural population. Given this, we chose to control the activity of the last hidden layer to best approximate the complexities of the brain.

**Figure 2.**
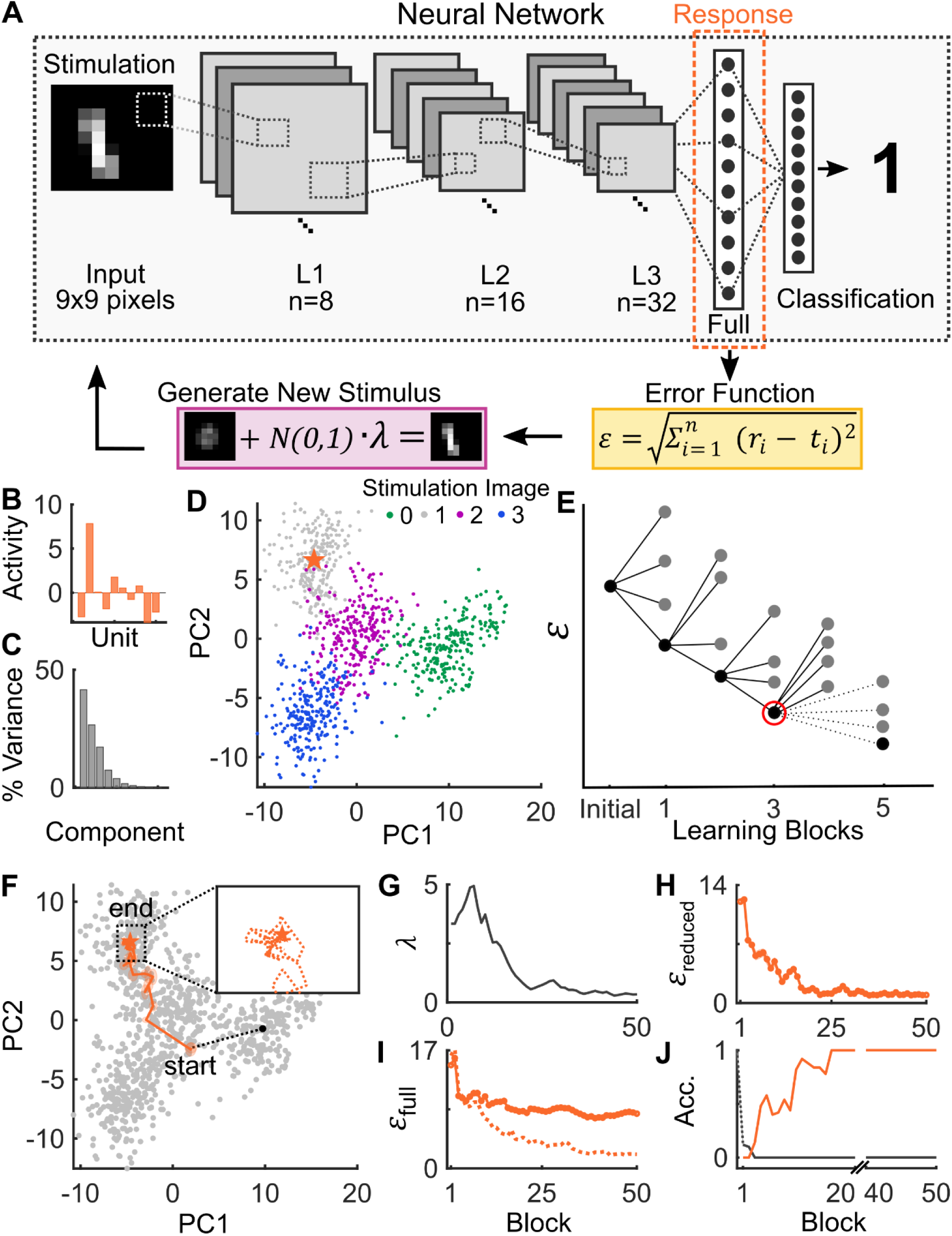
ACLS control of representations in a deep convolutional neural network. **(A)** Schematic of experiment. A 5-layer deep convolutional neural network (CNN) was trained to classify 10 numeric digits from the MNIST database (see Methods). Images were downsampled to a resolution of 9×9 pixels before training. Numerical images were delivered as ‘stimulation’ to the trained network and activity in the last hidden layer was considered the ‘response’. For each response, an error function (Euclidean distance) between the evoked and target response was computed (yellow box). This error was provided to the ACLS which used a stochastic learning algorithm (see panel E and Methods) to iteratively improve the stimulation pattern (purple box). **(B)** The target response of the CNN, taken as the response to a randomly selected image of the number 1. **(C)** Distribution of explained variance captured by the principal components (PCs) of the activity in the response layer of the CNN. **(D)** Response of the neurons in the response layer to 250 different images of the numbers 0, 1, 2 and 3, colored in green, gray, purple and blue, respectively. Responses are projected into the two-dimensional subspace created by the first two PCs. The orange star shows the target response (from B). **(E)** Schematic of ACLS stochastic learning algorithm. During a block, the algorithm generates a set of new patterns (column of dots) by perturbing the current ‘best’ pattern (black dot in previous column). Of these, the pattern that minimizes the error function of the system (y-axis) becomes the new ‘best’ pattern. This process repeats during each block, keeping the previous best stimulation pattern if none of the new patterns decreased error (red-circle and dotted lines). **(F)** Example learning trajectory of ACLS. Orange star denotes the target response. Orange line shows the trajectory of responses as the ACLS system learns. Each point denotes the mean response from a single block, with the shaded region around that point indicating the SEM in response (each block included 5 repetitions of 10 unique stimulation patterns). Initial orange point was mean response on block 1, with dotted black line indicating its displacement from the initial condition. In this simulation, Gaussian noise (µ = 0, σ = 0.5) was added to the response of the fully connected layer on each trial. Inset shows region indicated by gray box in full plot, with distributions of the ACLS learning path removed for clarity. **(G)** Annealing factor (λ) controls the magnitude of perturbations when generating new stimulation patterns. To arrive at global minima, while avoiding local minima, the stochastic learning algorithm increases/decreases the magnitude of the random perturbations to the current best pattern depending on whether a new best stimulus was/was not found during the previous block (see Methods and Box S1). **(H-I)** Error (ϵ) between ACLS-evoked responses and target response across blocks, computed in both the **(H)** 2D PC space and **(I)** full dimensional space. Solid line shows mean error per block (N=50 stimulations per block, x-axis). For comparison, the dotted line shows the error over learning when ACLS minimized error in the full-dimensional space (N=50 stimulations per block). **(J)** Stimulus classification accuracy by the CNN during learning. Orange line shows the fraction of ACLS-generated stimuli that were classified as the target category (e.g. the number 1). Black line shows the fraction of stimuli classified as the number 0 (the initial stimulation pattern provided to the ACLS; dotted line shows the classification rate of this initial image prior to the first block).

Our goal was to generate arbitrarily-defined patterns of activity in the last hidden layer. To this end, we began by mapping the response of this layer to 250 images of the numbers 0, 1, 2, and 3 (Fig. 2B; 1000 total images). As expected for a distributed representation, the first 2 principal components (PCs) captured the majority of the variance in activity of the target layer (68.14%; Fig. 2C). Similar low-dimensional representations are seen in the visual cortex, as shown below and in previous work (e.g. Cohen and Maunsell, 2010). Therefore, to reduce the impact of noise and to aid visualization, we projected the CNN response onto a two-dimensional space defined by the first two principal components (Fig. 2D, colored points).

A target response was randomly selected from the set of responses to images of the number ‘1’ (Fig. 2D, orange star). ACLS learned to produce this target response by minimizing an error function, taken as the Euclidean distance between the response of the CNN to the current stimulation pattern and the desired target response. Euclidean distance was calculated in the two-dimensional PC space (although similar results were observed in the full dimensional space, as shown below).

To learn to minimize this error function, ACLS used a stochastic learning algorithm (Fig. 2E). On each block of trials, the algorithm generated a set of new stimulation patterns by randomly perturbing the current ‘best’ stimulation pattern (see Methods). Each of these new patterns was then evaluated according to the error function and the stimulation pattern with the lowest error was taken as the new best stimulation pattern (if none of the new patterns improved performance, then the previous best stimulation pattern was kept). This process was repeated for each block, allowing the algorithm to iteratively move closer to a stimulation pattern that minimized the error function. In the example shown in Figure 2, ACLS was initialized with a random image of the number 0 from which to generate its first set of stimulation patterns. Note that the ACLS did not receive any information about the response to the images which generated the two-dimensional PC space or about the stimulation that produced the target response. The only input to the ACLS was the random initial stimulation pattern and the error of each stimulation response.

One concern with such ‘greedy’ algorithms is that they are prone to finding local minima in the error function. To reduce the likelihood of this happening, initial blocks of learning used large perturbations, allowing ACLS to broadly explore stimulation space. The magnitude of the perturbations was then slowly reduced over time, allowing the algorithm to identify a locally optimal stimulation pattern (see Methods and Box S1). The stochastic learning algorithm was chosen because it does not require a model of the system and because it does not have to estimate the gradient of the error function (avoiding strong assumptions about the nature of the response manifold). This is important, since neural responses may follow complex topologies in neural networks (DiCarlo et al., 2012). In this way, the stochastic learning algorithm avoids the obstacles that make it difficult to control complex neural networks.

As seen in Fig. 2F, ACLS successfully minimized the Euclidean distance between the stimulation-evoked neural response and the target pattern. As described above, the variance of the stimulation patterns was initially high, as the algorithm explored the broad stimulation space (Fig. 2G). Over time, the variance decreased, allowing the algorithm to settle on the (locally) optimal stimulation pattern. After 50 blocks of stimulation, ACLS discovered a stimulation pattern that was 92.32% closer to the target than its initial starting point in this example (Fig. 2H; 2500 total trials, 50 trials in each block, consisting of 5 repetitions of 10 stimulus patterns, see Methods). This was consistent across replications of the experiment (91.51 +/- 2.57% STD closer to target, N=50). Despite learning in a reduced-dimensional PC space, ACLS also reduced distance in the full-dimensional space, moving 48.25% closer in this space (Fig. 2I; 53.67 +/- 17.01% STD for N=50 replications). When ACLS minimized error in the full-dimensional space (rather than the 2D space), it moved 80.87% closer in this space (Fig. 2I, dotted orange line; 83.08 +/- 2.76% STD for N=50 replications). Interestingly, the stimulation patterns identified by ACLS were classified by the CNN as belonging to the target number (Fig. 2J). This categorization was not part of the error function but reflects the overlap between the stimulation evoked response and the original stimulus-driven response.

It is important to note that ACLS is not necessarily re-discovering the initial input that generated the target pattern (Fig. S1). Instead, ACLS is often discovering new stimulation patterns that produce the same response in the last hidden layer. Such ‘adversarial images’ are a known phenomenon in deep CNNs and reflect the convergent nature of the CNN (Szegedy et al., 2014). The goal of the CNN is to map varied inputs onto similar representations and so it is not surprising that the algorithm is identifying one of these alternative stimulation patterns. Indeed, if ACLS is tasked with controlling earlier hidden layers (e.g. convolution layer 1 or 2) the discovered stimulation pattern closely resembles the initial stimulation pattern (Fig. S1). This suggests the adversarial images are not a by-product of the ACLS approach but reflect the convergent nature of CNNs.

ACLS was robust to variability in neural responses. Previous work has shown neural responses are highly variable, possibly due to a combination of random noise and drift in the state of the animal (Calhoun et al., 2019). Here, we show ACLS is robust to both random noise and systematic drift. As seen in Figure 3A-B, ACLS was robust to random noise. Noise was modeled by adding white noise to the response to a stimulation pattern (Fig. 3A; see Methods). At low levels of noise (a signal to noise ratio, SNR, ≈ 18) ACLS learned at approximately the same speed as when there is no noise (Fig. 3B, purple and green lines, respectively). Intermediate levels of noise (SNR ≈ 9; yellow line) slowed the speed of learning, but ACLS still converges on a stimulation pattern that approximates the target response. Finally, high levels of noise (SNR < 4, gray and blue lines), disrupted learning significantly. This is to be expected, as the same stimulation led to drastically different responses from the CNN. In a high noise environment, averaging the responses across repeated stimulation helped to compensate for the impact of noise (Fig. 3B, light, medium, and dark gray lines show SNR ≈ 3 but with 1, 5, and 10 repetitions, respectively). This makes sense, as random noise averages to zero, but comes at the cost of slowing learning as each stimulation must be repeated. However, it suggests averaging the response to repeated stimulation can help ACLS learn in a high-noise environment, something we take advantage of when controlling neural representations *in vivo* (detailed below).

**Figure 3.**
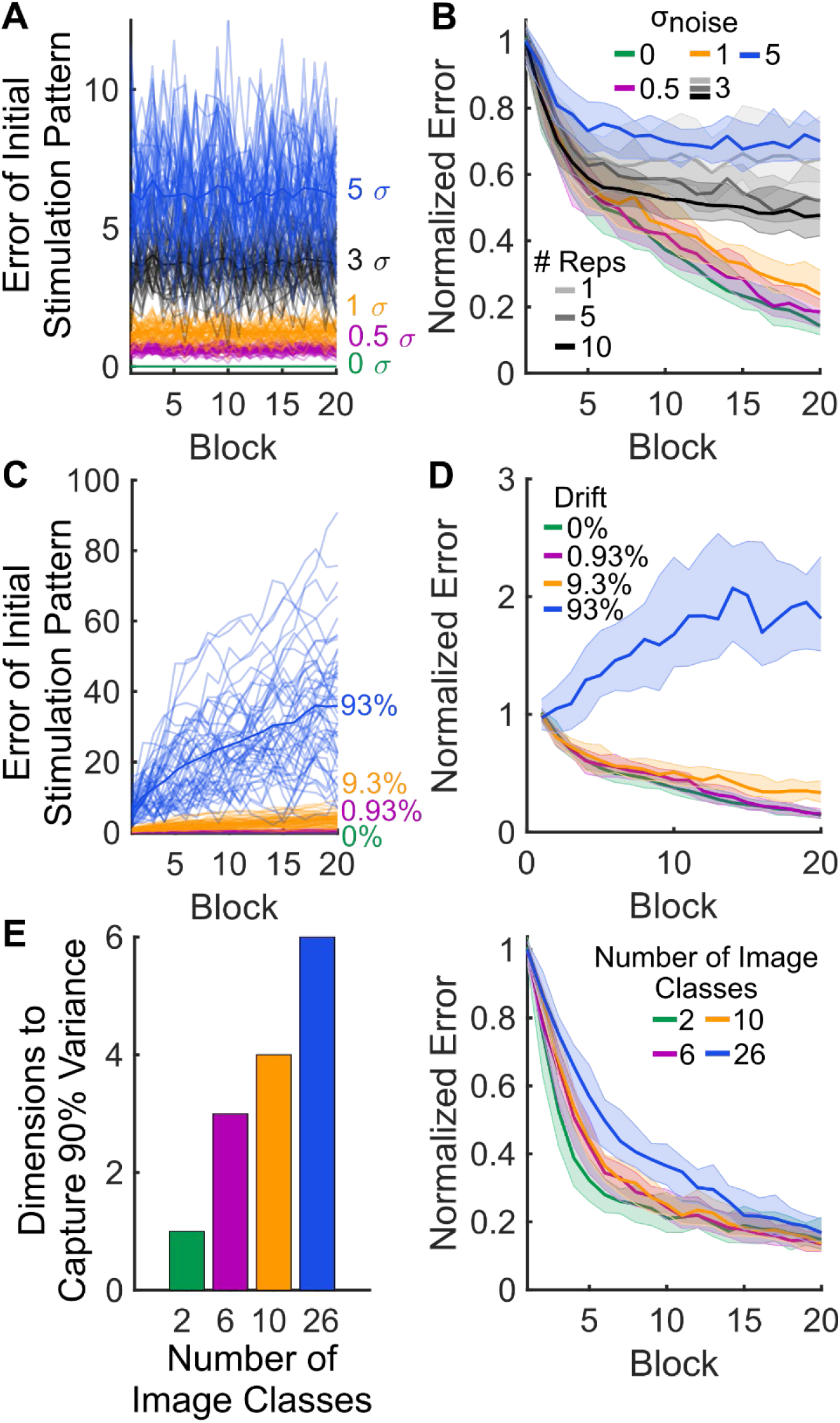
ACLS is robust to noise in the network, drift in the state of the network, and the complexity of the network. **(A)** Traces showing the impact of different noise magnitudes on the repeated presentation of the same stimulus pattern. On each trial, white Gaussian noise (µ = 0, σ = 0, 0.5, 1, 3, or 5) was added to the response of the final hidden layer (equates to an average SNR of ∼18, 9, 3, and 2, respectively). Traces show the block-averaged error of evoked response to the same stimulation pattern across blocks (e.g. the initial pattern given to ACLS; N=2 repetitions per block). N=50 replications for each noise level. **(B)** Noise in response impaired ACLS learning. Noise levels match those in A. Repeating stimuli mitigated the impact of noise: light, medium, and dark gray lines show the error across blocks with 1, 5, and 10 repetitions of each stimulus when noise was held constant at σ = 3. Lines show the median block-averaged error of evoked response, normalized to the average error of the first block (N=50 runs for each noise level). Shaded regions show inter-quartile range (IQR). **(C)** As in A, but shows how drift in the weights of the fully connected layer affects the response to the repeated presentation of the same stimulus pattern. On each trial, white Gaussian noise was accumulatively added to the weights of the fully connected layer (σ of noise distribution was 0, 0.93, 9.3, or 93% of the initial standard deviation in weights). N=50 replications for each drift level. **(D)** As in B, but shows the ACLS can compensate for most levels of drift. Colors match drift levels in C. **(E)** Bar plot showing that increasing the number of image classes to be classified by the CNN increased the dimensionality of the final hidden layer. Dimensionality was measured as the number of principal components needed capture *≥9*0% of the variance. **(F)** As in B and D, but shows increasing the complexity of the representational space slows ACLS learning, but does not prevent learning. Colors match image classes in E.

ACLS was also able to compensate for drift in the responses of the CNN. Drift was modeled by changing the weight matrix of the fully connected layer within the CNN (weights drifted by 0-93% per trial, see Methods). Changing the weight matrix meant the same input (i.e. an image of a ‘0’ or ‘1’) led to a different response over time (Fig. 3C). Despite this drift, ACLS was able to learn to generate the desired pattern, unless drift levels were such that the network was highly unstable (Fig. 3D). This is an advantage of the learning nature of ACLS; as the system state shifts, ACLS can learn to compensate for changes in the response.

Next, we investigated how the complexity of the CNN impacted the ability to learn. To manipulate complexity, we trained a new network to classify either 2, 6, 10, or 26 different letters (from EMNIST database, see Methods). Despite changes in the read-out layers, all other characteristics of the network remained the same. As seen in Figure 3E, increasing the number of stimuli to be classified increased the dimensionality of the hidden target layer (as measured by the number of PCs necessary to explain ≥90% of the variance in activity). In all cases, ACLS was able to learn to control the CNN. However, learning speed was slower in higher dimensional networks, suggesting more time may be needed to learn in complex networks (Fig. 3F; all ACLS parameters were held constant, see Methods).

Finally, as noted above, one of the strengths of the ACLS approach is that it can use different error functions to achieve different goals. To show this, we used ACLS to learn to control CNN activity with an error function that increased the cosine similarity between the evoked response and the target response (see Methods). Learning with this alternative error function was similar to previous results (Fig. S2A-B), suggesting ACLS can be used as a general tool for minimizing different error functions.

Altogether, our *in silico* simulations showed ACLS can learn to control a complex neural network. It was robust to noise, drifts in neural state, changes in network complexity, and could minimize multiple cost functions.

### ACLS control of neural representations *in vivo*

Next, we tested the ability of ACLS to control neural populations *in vivo*. ACLS was able to produce neural responses in both anesthetized and awake animals. Here we focus on awake animals as they provide the more complete test of ACLS; details of the anesthetized results are provided in the supplemental information (Figs. S3-4).

As with the *in silico* experiments, we were interested in testing whether ACLS could produce ‘natural’ neural responses. To this end, we began by mapping the response of visual cortex neurons to different visual stimuli (each presented for 200ms, see Methods and Fig. S5A). Both electrophysiological recording and electrical stimulation were done through a single silicon probe inserted into primary visual cortex (V1; 32 channels for recording, 32 channels for stimulation, see Figs. 4A and S5B-C and Methods). This allowed us to record from small populations of neurons (5-11, all multi-unit activity), while stimulating at nearby sites. The response of a multi-unit neuron to a visual stimulus was taken as the number of spikes during a 200ms window after stimulus onset. The precise timing of the window varied across recordings to best capture the evoked response but was typically 40-240ms after stimulus offset (see Methods). Multi-units that were not selectively responsive to the visual stimuli were excluded from further analysis, including ACLS learning (see Methods for exclusion criteria). The response of selective neurons was then used to define a population vector response to each stimulus.

**Figure 4.**
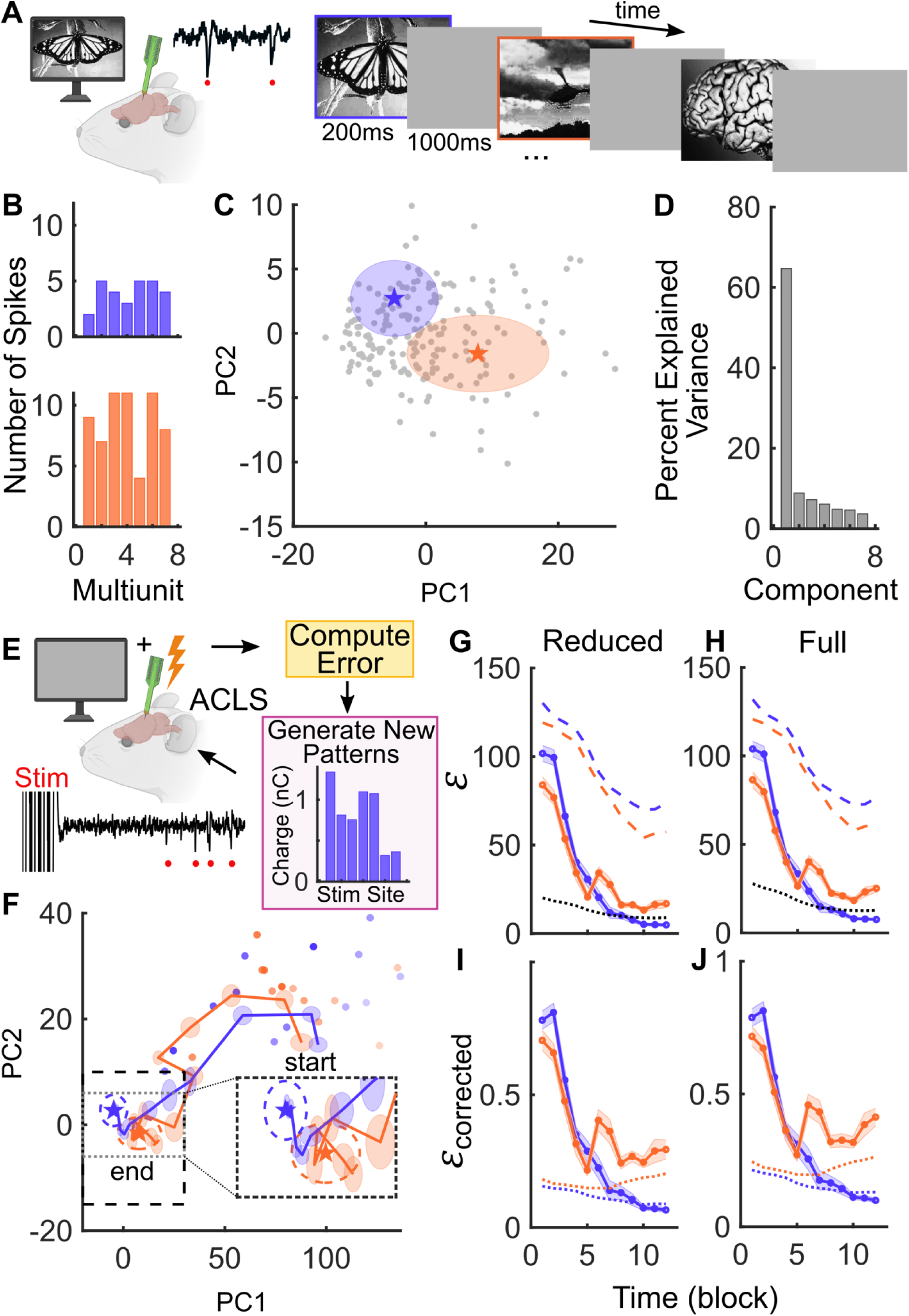
ACLS can control neural representations in vivo, in the primary visual cortex of awake mice. **(A)** Experiment schematic. Awake, head-fixed mice were presented with visual stimuli while recording multi-unit neural activity in the primary visual cortex (V1). Visual stimuli were 200ms in duration, interleaved with 1000ms presentations of a gray screen. Black trace shows example raw trace of visual stimulus-evoked multi-unit spiking activity (red dots). **(B)** Example multi-unit activity response to the purple (top) and orange (bottom) outlined images in A. **(C)** Distribution of responses to 10 different visual stimuli, each presented 20 times (gray dots indicate response to one stimulus). Purple and orange stars show the target response to the purple and orange-outlined stimuli. Shaded regions denote STD of 20 visual stimulus presentations of each stimulus. **(D)** Explained variance in multi-unit activity captured by the principal components. **(E)** Schematic of ACLS learning to reproduce visual stimulus-evoked neural responses. Electrical stimulation was applied with a gray screen to match baseline visual input. An example stimulation pattern across 7 stimulation sites is shown in purple. Black trace shows example raw trace of stimulation followed by multi-unit spiking activity (red dots). **(F)** Example timecourse of learning. Orange and purple stars indicate the two target responses (as in B and C, dotted lines around each star shows STD of visual response). Targets were learned by two simultaneous, but independent, runs of the ACLS algorithm. Orange and purple lines show the 2D trajectory of learning over blocks. Shaded region around each step indicates the SEM of responses in that block (N=12 blocks, each with 5 repetitions of 6 unique stimulation patterns). Individual purple and orange points show response to repetitions of the initial stimulus pattern (1 repetition per block). Transparency of points scales with time (light → dark). The movement of these points show the system drifts over time. Black dotted line denotes borders of 2D visual response space shown in C. Inset shows region denoted by gray box in full sized image. Note that the only input to the ACLS algorithm was the initial stimulation pattern (purple pattern in E) and the error between each evoked response and the target response (purple and orange lines). **(G-H)** Error between ACLS-evoked responses and target response across blocks in **(G)** 2D response space and **(H)** full-dimensional space. Solid orange and purple lines show mean error per block (Euclidean distance; N=30 stimulations); shaded regions show SEM per block. Dotted black line denotes noise floor, quantified as the average pairwise Euclidean distance between repetitions of the same stimulation pattern (N=5 repetitions of 6 unique stimulation patterns per block). Dashed purple and orange lines show the drift in the response of the initial stimulation, measured as the distance to the target response. Lines for initial stimulus repetitions and noise lines reflect sliding window mean of 6 blocks. **(I-**Error over time, corrected for drift of initial stimulation response over blocks (see Methods). As in G-H, solid lines show error of learned responses across blocks. Dashed lines show the corresponding noise floors (also corrected).

Figure 4A shows an example recording in which an animal received 20 repetitions of 10 different isoluminant visual stimuli, resulting in 200 stimulus responses (Fig. 4B, response vector is across 7 multi-units). To define the response space of the recorded visual cortex neurons, we used PCA to create a two-dimensional space that captured the majority of variance in responses (73.53%, Fig. 4C-D; across N=24 recordings the first 2 PCs captured 81.54% +/- 1.77% SEM of the variance). Projecting responses into this reduced dimensionality space mitigated the effect of noise and emphasized the effect of visual stimulation. It also allowed us to define the space of ‘achievable’ patterns of neural responses.

Once the visual response space was defined, we used ACLS to learn electrical stimulation patterns that could reproduce the neural response to visual stimuli. Electrical stimulation was delivered through 32 dedicated stimulation electrodes (see Fig. S5B for example map of stimulation electrodes on the neural probe). Although 32 stimulation channels were available, several of them fell outside of cortical tissue. So, stimulation was limited to sites that were local to recording electrodes with a neural response and, therefore, were likely in or near cortex (typically 6-16 stimulation sites; see Methods for exclusion criteria). Electrical stimulation patterns were defined as the vector of stimulation amplitude across the set of stimulating electrodes. Each electrode delivered a specific amount of charge (between 0-10 nC), that was updated by the ACLS system from trial to trial. All stimulations were delivered in a sequence of 10 cathode-leading, high-frequency bi-phasic pulses (200µs pulses delivered at 300 Hz). The neural response to electrical stimulation was measured in the same window as visual stimulation (e.g. 200ms window starting 40-240ms post stimulation offset), which avoided artifacts from the electrical stimulation (see Fig. S5C for schematic of spike detection and Methods).

Before learning, two target responses were manually selected from the visual stimulus response space (Fig. 4C, orange and purple stars). Selection was done blind to the exact response pattern, but targets were chosen such that they were separated in the neural response space. The ACLS algorithm then learned to produce each target response by reducing the error between the evoked response and the target (see Fig. S6 for example stimulations and responses along the learning trajectory). As for *in silico* experiments, the error function was taken as the Euclidean distance in two-dimensional PC space between the evoked response and the target response. The ACLS system had no *a priori* knowledge of the stimulation space or the relationship between stimulation and responses. ACLS was only provided with *a)* a randomly-generated initial stimulation pattern (Fig. 4E; each channel was randomly drawn from a uniform distribution, bounded to prevent tissue damage), and *b)* the error of each evoked response. The two selected targets were learned simultaneously by two independent runs of the ACLS algorithm (alternating randomly between the two targets; no information was shared between runs).

As with the *in silico* experiments, ACLS learned to produce the target pattern of neural responses. Figure 4F shows ACLS learning during our example session (see Supplemental Information and Fig. S7 for additional examples). Across 12 blocks, the evoked responses systematically moved through the two-dimensional PC space, moving 79.90% and 95.20% closer to their respective targets (i.e. orange towards orange and purple towards purple; 30 trials are in block, consisting of 5 repetitions of 6 stimulus patterns, see Methods for details). Accordingly, the error between the evoked response and the target response decreased over time (Fig. 4G; orange and purple solid lines, shading denotes SEM). Although the system was attempting to minimize error in the two-dimensional PC subspace, error was also reduced in the full dimensional neural space (Fig. 4H). Stimulation occurred approximately every 2.1 seconds, resulting in a total learning duration of ∼12.5 minutes for each run (∼25 minutes in total for the 2 simultaneous runs).

As noted in our analysis of the *in silico* model, both noise and drift in neural responses influence the ability of ACLS to learn to evoke a desired response. Here, noise was estimated by the average pairwise distance between the neural response to repetitions of the same stimulation pattern. This estimate of noise is shown in Figure 4G-H (black line). Such noise can impact ACLS in two different ways. First, it can lead to noise in the estimate of the error of a particular stimulation pattern (as shown *in* silico, Fig. 3A). To compensate for this, ACLS averaged over 5 repetitions of each stimulation pattern. Second, the noise in the response also places a fundamental limit on how close ACLS can get to the desired response pattern – it is impossible to get closer than the ‘noise floor’. This can be seen in Figure 4G-H: the error between the evoked response and the target response reduces across blocks but asymptotes close to the noise floor (Fig. 4G-H, black dashed line).

Drift in the system is reflected in a systematic change in the response to the same stimulation over time. Such changes could reflect adaptation to stimulation, movement of the electrode, or changes in the animal’s physiological state (i.e. arousal). To understand the nature of drift *in vivo*, we repeated a fixed set of 10 stimulation patterns in an anesthetized mouse over a period of 1 hour. As seen in Figure S8, the response to the same stimulation pattern changed dramatically over time, suggesting drift can be a significant issue. Interestingly, the magnitude of drift was highly correlated across stimulation patterns, suggesting drift reflects a global state change, rather than pattern-specific changes.

It is important to account for drift when evaluating the success of ACLS, as drift could either be towards or away from a particular target. Therefore, to measure and correct for drift, we repeatedly delivered the initial stimulation pattern for both targets throughout the recording (note: the ACLS algorithm was not aware of these stimulations or the responses). In our example recording, the evoked response to the initial stimulation patterns drifted over time (Fig. 4F, orange and purple dots), which decreased the ‘error’ of the initial stimulation pattern relative to the target responses (Fig. 4G-H, orange and purple dashed lines). To correct for this drift, we calculated the ‘drift-corrected error’ of each response by dividing the error of each response by the ‘error’ of repeats of the initial stimulation pattern (see Methods). As seen in Figure 4I-J, even following this correction, ACLS is able to reduce the error between the evoked and target response in both the reduced and full dimensional spaces.

A decrease in drift-corrected error was observed for the majority of recordings. Figure 5A shows the error traces of all ACLS recordings (48 ACLS runs, across 24 recordings in 9 animals), colored by whether ACLS successfully reduced error. Across all successful runs, ACLS decreased the Euclidian distance by 36.91% +/- 4.90% SEM by the end of the 5^th^ block and by 45.57% +/- 3.75% SEM by the end of the 10^th^ block (often approaching the noise floor, Fig. 5B). To quantify learning of each run, we measured the difference in error between the evoked and target response on the first and last block of learning. Over 48 total runs, this error was reduced in 85.42% of the runs (Fig. 5C; 41/48 runs, p=10^−7^, binomial test vs 50% chance). Similarly, we can quantify the change in error with learning by fitting an exponential curve to the error over the blocks in each run (*f(x) = ae*^*λx*^ + *c*, see Methods). Using this metric, 81.25% of the runs successfully reduced error (λ<0 for 39/48 runs, p=10^−6^, binomial test). In the full dimensional space, ACLS reduced error on 81.25% of the runs (quantified as decreased error in the last block compared to the first; 39/48, p<=10^−6^, binomial test).

**Figure 5.**
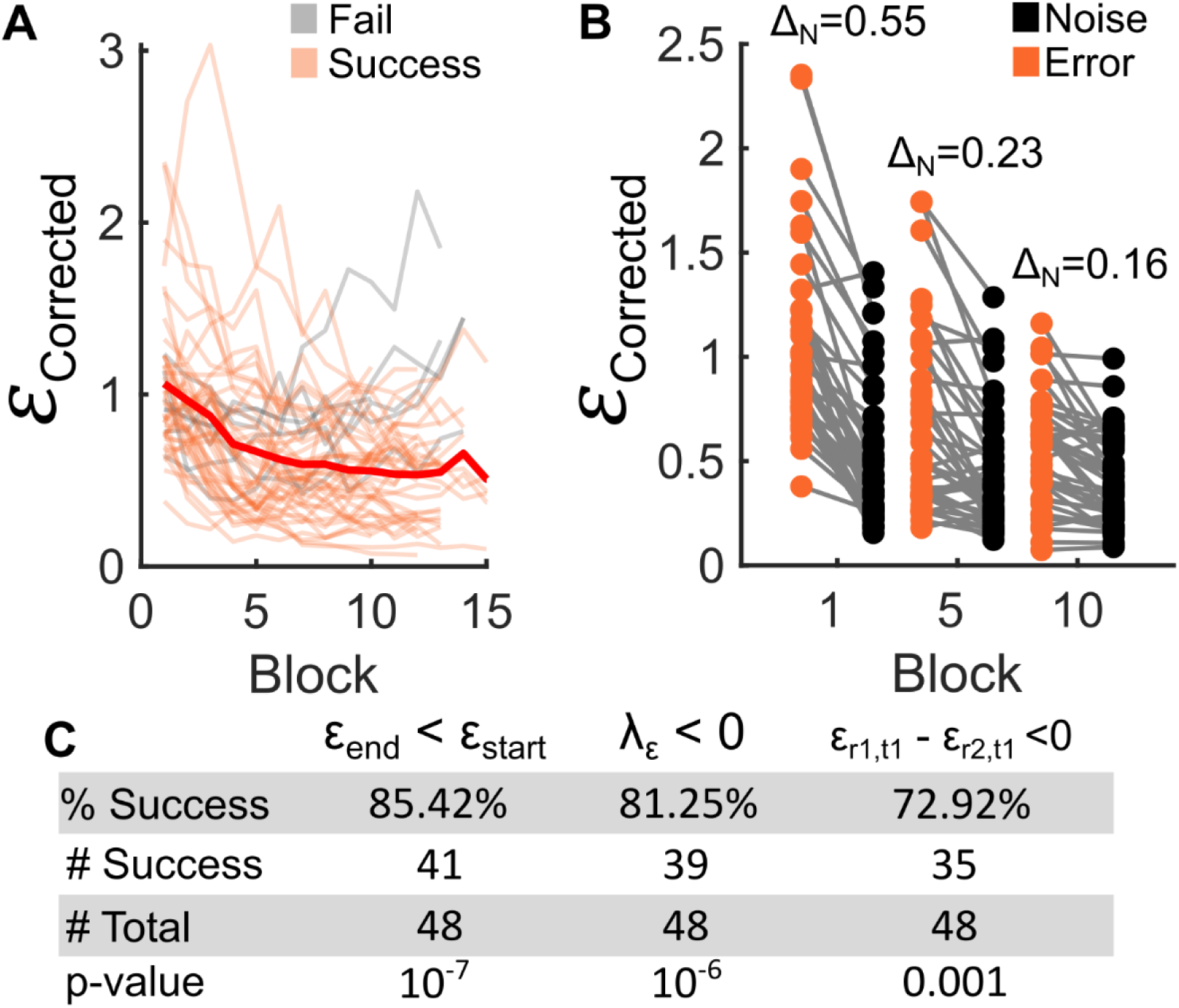
Quantifying ACLS success in awake animals across multiple sessions. **(A)** Corrected error over time for all ACLS runs (N = 48) in all recording sessions (N=24), across all mice (N=16). Display follows Figure 4I. Individual orange and gray lines show error over time of individual successful and failed ACLS runs, respectively. Runs were considered successful if the mean error during the final block was less than the first block. Red line shows mean trajectory across successful runs. For visualization, the x-axis truncated at 15 blocks (N=6 runs were longer than 15 blocks, all continued to decrease in error). **(B)** Pairwise comparisons of drift-corrected error (orange dots) and noise (black dots) at blocks 1, 5 and 10 across all ACLS runs (N=48). Δ_N_ is the mean difference between the error and the noise of each run (i.e. gray lines). **(C)** Table of success rate of the ACLS runs across multiple sessions, quantified by a reduction in error over learning (left column), a negative exponential fit to the learning curve (middle column), and specific learning towards one target (relative to the other; right column). See text and Methods for details of the statistics.

One concern is that these success metrics may reflect ACLS learning to navigate from the initial stimulation pattern to the ‘cloud’ of visually-evoked responses, rather than learning to produce a specific pattern of firing activity. To test the ability of ACLS to selectively evoke population-level neural responses, we calculated whether a given ACLS run ended closer to its target than the other simultaneous run (e.g. *ε*_*run*1,*target*1_ − *ε*_*run*2,*target*1_ < 0). Consistent with selective responses, 72.92% of runs were selective for their own target in both the reduced and full dimensional space (Fig. 5C; 35/48 runs, p=0.001 by binomial test for both spaces). As an alternative metric, we calculated whether a given run ended closer to its target than the other target (e.g. *ε*_*run*1,*target*1_ − *ε*_*run*2,*target*2_ < 0; drift-corrected to compensate for a global shift towards either target). Using this metric, 68.75% of runs were selective for their own target in both the reduced and full dimensional space (33/48, p=0.0066 by binomial test for both spaces). One may expect selectivity values to be lower than overall success rates since they are more susceptible to high noise levels blurring discriminability between the targets. Indeed, logistic regression found the signal to noise ratio of the run was predictive of selectivity (p=0.018, see Methods for details). Supplemental experiments further validated the selectivity of ACLS in anesthetized experiments; it could selectively navigate between two arbitrarily-defined, electrically-evoked neural responses (Figs. S3 and S4; see Supporting Information).

### Using Adaptation to Test the Physiological Relevance of Electrically Evoked Patterns

As noted above, our goal was to test whether ACLS could produce ‘natural’ neural responses. So far, we have measured similarity between the visually-evoked and electrically-evoked as the Euclidean distance in neural space. One concern is that this may result in neural responses that are not physiologically meaningful, either because Euclidean distance may not be an accurate measure of response similarity or because learning occurred in a reduced dimensional space (even though the full N-dimensional error was also reduced).

Therefore, to test the physiological similarity between electrically-evoked and visually-evoked responses, we tested whether electrically-evoked responses adapted to visual stimuli. Previous work has shown that the neural response to a visual stimulus is reduced when the stimulus is repeated (Jin et al., 2019). This ‘adaptation’ of the neural response has been used to measure the similarity of neural responses to visual stimuli (Leopold et al., 2001). Here, we used this approach to test if neural responses to electrical stimulation were physiologically similar to visually-evoked responses by testing if the response to electrical stimulation was adapted by the presentation of the associated visual stimulus.

To this end, we first mapped the visual response of neurons and then used ACLS to learn stimulation patterns that generated neural responses similar to two different visual stimuli (as in Fig. 4 and 5). Specifically, electrical stimulation pattern ES1 and ES2 were taken as the stimulation patterns with the smallest error to visual stimuli VS1 and VS2, respectively (from last block of learning). To ensure the ACLS was able to reproduce the visual response, we only included the experiments in which ACLS successfully decreased the error towards both targets (N=4 total sessions).

Next, we tested the effect of adaptation on both visually-evoked and electrically-evoked responses. Animals were presented with a sequence of three stimuli; two visual stimuli, followed by an electrical stimulation (Fig. 6A; see Methods). The sequence always started with visual stimulus 1 (VS1) or visual stimulus 2 (VS2). This was followed by either the same stimulus (e.g. VS1→VS1) or the other stimulus (e.g. VS2→VS1). Finally, the electrical stimulation could either match the second visual stimulus (e.g. VS1/2→VS1→ES1) or be different (e.g. VS1/2→VS2→ES1). Figure 6B shows the effect of adaptation for two example neurons. As expected, repeated presentation of the same visual stimulus led to adaptation, reflected by a reduction in firing rate to the repeated stimulus (neuron 1: 15.16% in 100ms-150ms and 9.47% in 150ms-200ms, Fig. 6B, upper-left; neuron 2: 7.94% in 100ms-150ms and 9.83% in 150ms-200ms, Fig. 6B, bottom-left). The response to ACLS-learned electrical stimulation patterns was also adapted when preceded by the matching visual stimulus (neuron 1: 4.79% in 100ms-150ms and 0.84% in 150ms-200ms, Fig. 6B, upper-right; neuron 2: 20.13% in 100ms-150ms and 54.21% in 150ms-200ms, Fig. 6B, bottom-right; see Methods).

**Figure 6.**
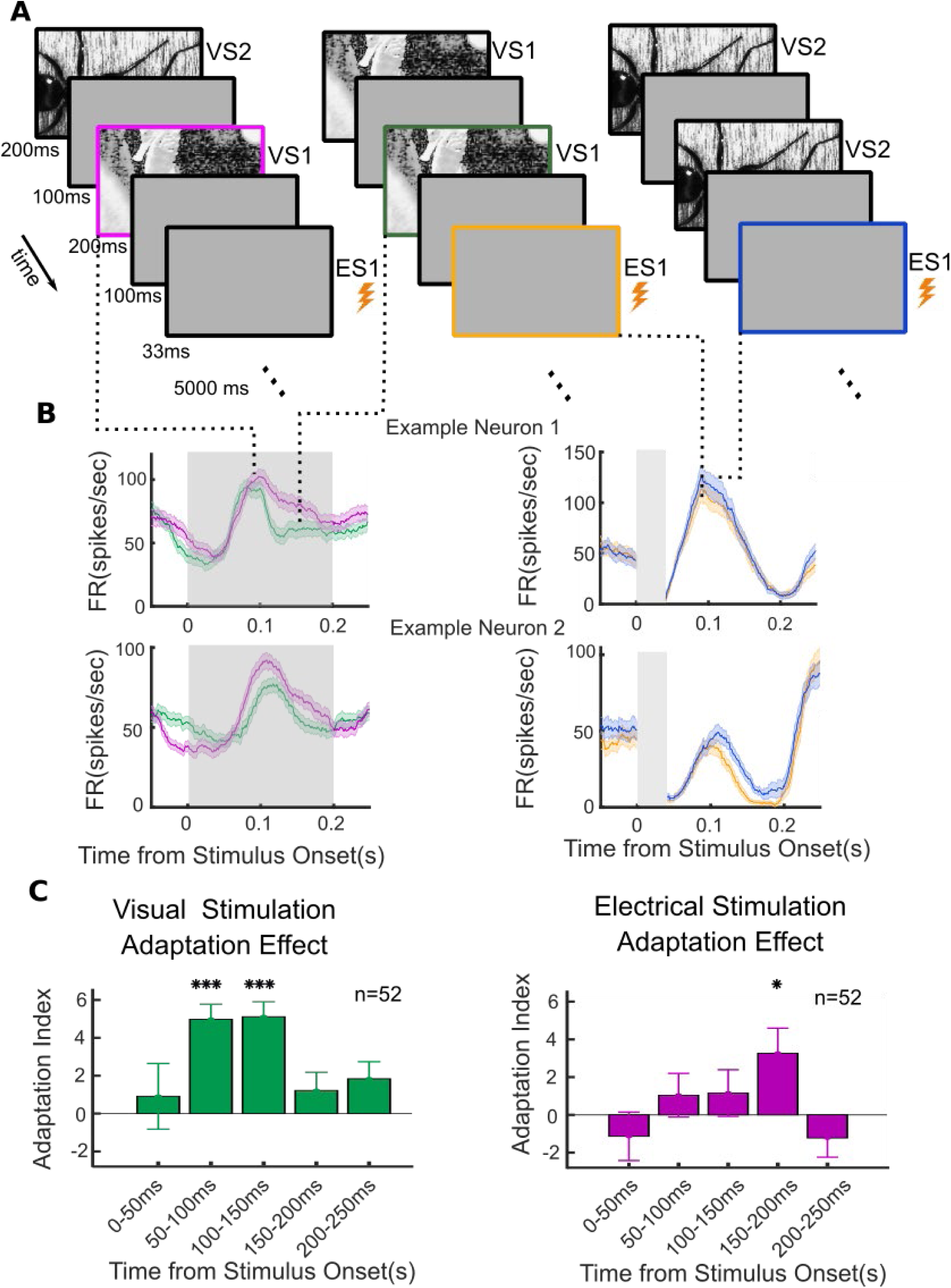
ACLS-learned electrical stimulation elicits responses that are physiologically similar to the response to visual stimuli. **(A)** Schematic of the adaptation paradigm. On each trial two visual stimuli (VS1 or VS2) were followed by an electrical stimulation (ES1 or ES2). This yielded 8 total conditions, 3 of which are shown here (see Methods for full list of conditions). Visual stimuli were 200ms in duration, interleaved with 100ms presentations of a gray screen. ES1/ES2 were the stimulation pattern with the smallest error to the response evoked by VS1/VS2. A 4.5-5s inter-trial interval (ITI) separate each trial. **(B)** Two examples of adaptation in multi-unit activity (one example per row). Left plot shows the response to VS1 is lower when preceded by VS1 (green line) than when preceded by VS2 (purple line). Traces show peristimulus time histogram, smoothed with a 50ms moving average. Right plot shows the response to ES1 was lower when it was preceded by VS1 (yellow line) compared to when it was preceded by VS2 (blue line). Gray regions indicate stimulus presentation period. **(C)** Adaptation was a consistent effect across all recorded neurons. The magnitude of adaptation during visual (left) or electrical (right) stimulation. Adaptation was measured by an adaptation index, calculated as the percent change in firing rate in response to a different (or non-associated) stimulus (e.g. VS1*→* ES2) relative to a repetition of the same (or associated) stimulus (e.g. VS1*→* ES1; see Methods). Bar plots show mean and SEM of adaptation index in 50ms sliding windows across all stimulus pairs and all recorded multi-units (N=52 pairs from N=4 sessions; ***p_bonferroni_<0.001, *p_bonferroni_ <0.05)

Adaptation was consistently observed across sessions and neurons (Fig. 6C; N=4 sessions, N=26 neurons). The evoked response to a repeated visual stimulus was reduced by 5.04% in the 50-150ms following stimulus onset (p<=5.5*10^−8^ in 50-100ms and p<= 5.6*10^−8^ in 100-150ms, t-test with Bonferroni-corrected p-value, Fig. 6C). Similarly, the response to electrical stimulation was reduced by 3.26% in the 150-200ms following stimulus onset if it was preceded by the matching visual stimulus (p=0.0092, t-test with Bonferroni-corrected p-value). Note that this effect followed the time course of the response to electrical stimulation (and not visual stimulation). This is to be expected, as adaptation is modulating the response to visual/electrical stimulation, not reshaping its time course. Together, these results suggest the neural responses evoked by ACLS are physiologically similar to natural neural responses.

## Discussion

Here we present an Adaptive Closed-Loop Stimulation (ACLS) system that can learn to produce arbitrarily-defined neural responses. The system uses a stochastic learning algorithm to reduce the error between the evoked neural response and a target neural response. Importantly, the learning algorithm allows the ACLS system to be agnostic to the underlying structure of the neural circuit and avoid having to model how neurons respond to electrical stimulation.

Here, we show ACLS works both in simulated and real neural networks. ACLS learns to generate arbitrarily-defined neural responses that match the response to visual stimuli (Fig. 4, 5 and 6). In awake animals, learning was successful for the majority of the recording sessions (∼80% of runs reduced their error). The error between the initial and target pattern was typically reduced by ∼50% (often approaching the noise floor). This reduction was done quickly, within ∼15 minutes (∼10 blocks with ∼300 total stimulations). In addition to reducing the error in ‘neural space’, we found electrically evoked responses were physiologically similar to natural responses evoked by visual stimuli. This was seen in the adaptation of electrically-evoked responses to a preceding, matching visual stimulus. Importantly, ACLS was robust to noise and drift in the state, as shown in both artificial and real neural networks.

Here we present the ACLS using electrophysiological recordings of neural activity and electrical stimulation. However, the ACLS framework is flexible. It can be easily adapted to work with a variety of stimulation approaches (e.g. electrical, optical, or magnetic), a variety of ways to measure the response of the brain (e.g. single units, bold signal, local field potentials, or behavior), and a variety of error functions to achieve a desired neural or behavioral state. We believe this flexibility will allow ACLS to have several unique scientific and clinical applications.

### Scientific Applications of ACLS

Compared to standard neural stimulation techniques, ACLS allows for 1) improved control of neural populations and 2) optimization of arbitrary error functions. Both of these characteristics will enable new scientific applications.

Recent experimental work has shown information in the brain is represented in a distributed manner, across the neural population (Saxena and Cunningham, 2019; Yuste, 2015). These population responses are thought to reside on a low dimensional manifold (Cunningham and Yu, 2014), with the position of neural activity on this manifold representing the value of a cognitive variable (e.g. visual perception, DiCarlo and Cox, 2007). However, this hypothesis has been difficult to causally test because, in part, it requires the ability to control a population of neurons. The ACLS could address this gap by learning stimulation patterns that evoke arbitrary neural responses along the manifold. This would allow one to test how cognitive variables evolve over the manifold (as suggested by Jazayeri and Afraz, 2017). In particular, because ACLS can learn arbitrarily-defined neural responses, one could directly compare the behavioral response to neural activity that is either on or off the manifold (Sadtler et al., 2014; Wärnberg and Kumar, 2019).

In addition to controlling populations of neurons, the ACLS approach may also be able to change other characteristics of neural activity. This would simply require changing the error function. For example, ACLS could potentially learn to induce an oscillation within or between neural populations, allowing one to directly test the predicted role of oscillations in cognition (Danzl et al., 2009; Fries, 2015; Helfrich et al., 2018; Knudsen and Wallis, 2020).

### Clinical Applications of ACLS

Clinically, ACLS may improve the precision and efficacy of clinical neuromodulation devices. By learning the appropriate stimulation patterns, the system will be able to optimize stimulation protocols for neurodegenerative and neuropsychiatric disorders. In particular, by defining a disease-specific error function, ACLS could optimize the stimulation pattern to reduce a specific neural or clinical symptom. For example, one might want to treat Parkinson’s disease by stimulating such that pathological beta-band oscillations are reduced (Little et al., 2013). Alternatively, one could stimulate in a way that reduces motor tremors (Groiss et al., 2009). In both cases, the same ACLS approach is used; only the error function is updated. Furthermore, because error functions can be combined, the best stimulation pattern may be one that minimizes both beta-band oscillations and tremors. In this way, ACLS could provide the ability to extend existing stimulation treatments to allow for greater flexibility in the stimulation pattern and treatment options.

Furthermore, the adaptability of ACLS could help to individualize treatments. Recent work suggests that the efficacy of deep-brain stimulation depends on physiological differences between patients and the exact placement of stimulating electrodes (Greene et al., 2020; Horn, 2019). The adaptability of ACLS could allow for a greater number of electrodes to be implanted, with the system searching for the optimal stimulation pattern across those electrodes. Furthermore, current neurostimulators cannot quickly adapt to changes in state (i.e. changes in wakefulness or progression of a disease). Again, the on-line, adaptive, nature of ACLS will allow it to automatically compensate for such state changes by continuously tracking its own performance.

### Future directions

In its current form, ACLS has several limitations that motivate future work. Here, we used a simple stochastic learning algorithm. Although we show that this approach can be successful, more sophisticated optimization algorithms may improve the success rate and speed of learning. Similarly, the real-time nature of learning limited us to using a simple threshold detection to identify multi-unit neural activity. This likely increased the level of noise in recorded responses and limited the selectivity of ACLS learning. Thus, real-time spike sorting may improve future ACLS performance. Future work is also needed to test the ability of ACLS to learn over longer periods, while controlling more complex stimulation systems (greater than the ∼10 stimulation sites used here) with more precise temporal control (beyond the 200ms time window used here). Finally, future work is needed to test the ability of ACLS to improve clinical outcomes in patients.

## Acknowledgments

The authors thank Alex Libby, Caroline Jahn, Matt Panichello, and Sarah Henrickson for their feedback. We also thank the Princeton Laboratory Animal Resources staff for their support. This work was funded by NIH DP2 EY025446.

## Author Contributions

Conceptualization: TJB, ST, CJM, KCL; Data curation: ST, CJM, TJB; Formal Analysis; ST, CJM, TJB; Funding acquisition: TJB; Investigation: ST, CJM, DC, KCL, TJB; Methodology: ST, CJM, TJB; Project administration: TJB; Resources: TJB; Software; ST, CJM, TJB, KCL, CS; Supervision: TJB; Validation; CJM, ST; Visualization: CJM, ST, TJB; Writing – original draft: ST, TJB, CJM, DC; Writing – review & editing: TJB, ST, CJM, DC, KCL, CS.

## Methods and Supplemental Information

Key Resources Table

| REAGENT or RESOURCE | SOURCE | IDENTIFIER |
| --- | --- | --- |
| Experimental Models: Organisms/Strains |  |  |
| Mouse: C57Bl/6J, GP5.3, PV<sup>Cre</sup> | The Jackson Laboratory | #000664, 028280, and 008069 |
| Software and Algorithms |  |  |
| MATLAB 2017a-2019a | Mathworks | https://www.mathworks.com/products/matlab.html |
| Adaptive Closed Loop Stimulation Package | This paper | https://github.com/buschman-lab |

### Lead Contact and Material Availability

Request for additional information, resources, and reagents should be addressed to Lead Contact, Timothy J. Buschman.

### Experimental Model and Subject Details

All experiments and procedures were carried out in accordance with the standards of the Princeton University Animal Care and Use Committee (IACUC) and the National Institute of Health. Adult female (N=7) and male (N=2) mice of at least 12 weeks of age were used for awake experiments and male (N=5) and female (N=5) mice were used for anesthetized experiments (strains used: C57Bl/6J, GP5.3 heterozygotes, or PV^Cre^ heterozygotes; Jax Stock numbers 000664, 028280, and 008069, respectively). Animals were single housed on a reverse 12-hr light cycle and received standard rodent diets and *ad libitum* water. All experiments were performed during the dark period, typically between 12:00 and 19:00. A subset of the mice (N=4) used for awake experiments were also used for adaptation experiments.

### Methods

### Adaptive Closed-Loop Stimulation System Implementation

The Adaptive Closed-Loop Stimulation (ACLS) system was implemented in a custom-built MATLAB software package and GUI. Pseudocode outlining ACLS framework is shown in Box S1. The GUI enabled users to define experimental parameters, view neural stimulation and response in real time, provide visual stimuli, and batch run simulations. For *in vivo* experiments, the GUI interfaced with a microstimulation system (RZ5 BioAmp Processor, Tucker Davis Technologies) to deliver electrical stimulation and record neural response in real time.

The first component of ACLS is observing the evoked response to a specific pattern of stimulation. For *in silico* experiments, the evoked response was the activity of one of the CNN hidden layers (detailed below). For *in vivo* experiments, the neural response was the number of spikes from recorded multi-units during a predefined window (detailed below).

The second component of ACLS is evaluating an error function to determine the proximity of current algorithm state to the target response. For all *in vivo* experiments, the error function was defined as the Euclidean distance between the evoked response (*r*) and the target response (*t*):

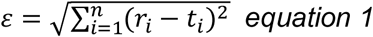

where *r* and *t* are the coordinates in the *n*-dimensional space (e.g. either 2D PC or full ND space, where N is the number of recorded multi-units). Importantly, ACLS is compatible with any error function (e.g. Euclidean distance, correlation, cosine similarity; see Fig. S2A-B for example of ACLS learning with a cosine similarity error function).

The third component of ACLS is to use a machine learning algorithm to automatically update stimulation parameters to reduce the error function on the subsequent iteration. Here, this was achieved using a ‘greedy’ stochastic learning algorithm. This algorithm generated new stimulation patterns for each iteration by randomly selecting from a distribution around the previous best stimulation pattern (gaussian distribution for *in silico*, uniform bounded distribution for *in vivo* experiments). The previous ‘best’ stimulation pattern was the stimulation pattern that evoked the lowest error (averaged across repetitions) during the previous 4 blocks.

To find global minima while avoiding local minima, the spread of this distribution was determined by an ‘annealing factor’, λ. The value of λ changed after each block, according to:

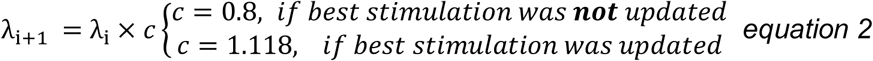

where λ_*i*_ and λ_*i*+1_ are the current and next λ values respectively, and *c* is a weighting scalar. If a new, ‘best’ (e.g. minimum error), stimulation pattern was found on the current block, then *c =* 1.118 to increase the search space. If no new best was found, then *c =* 0.8 to settle to the minima. λ was initialized at 3.3 and capped at 10 for *in silico* and *in vivo* recordings. For example, given Eq. 2, the new set of *n* stimulation patterns for *in silico* experiments would be:

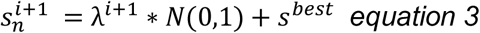

where *s*^*best*^ is the previous best stimulation pattern and 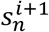 are *n* new stimulation patterns.

ACLS has several parameters, including the number of stimulation patterns generated in each block, the number of repetitions of each stimulation pattern, and the scaling of the annealing factor with learning (*c* from Eq. 2). In the main and supplemental text, we explored different values for these parameters (Fig. 3; Fig. S2). Based on this, we found 5-10 new stimulation patterns per block balanced the speed of learning with the degree of exploration in stimulation space (Fig. S2C). Increasing the number of stimuli increased the block length, which decreased the rate of learning. This could be problematic if the features of the system being controlled change more rapidly than the algorithm can learn (i.e. it would no longer be able to compensate for drift). However, if the system is stable, then more stimulation patterns will improve the ability of ACLS to explore stimulation space and reliably find a globally optimal stimulation pattern (seen as decreased variance in learning curves when more patterns are used in each block, Fig. S2C). As shown in Figure 3, the number of repetitions of each stimulus should be chosen to mitigate the impact of noise while avoiding unnecessary resampling (that will slow overall learning). We found averaging over 3-5 repetitions worked well *in vivo*. We did not systematically explore different scaling factors of the annealing (*c* in Eq. 2). However, based on sparse sampling, we found setting *c* to [0.8, 1.118] was reasonable and allowed for exploration while still settling to a minimum in a reasonable amount of time. Increasing *c* will broaden exploration, while decreasing *c* will minimize exploration and allow ACLS to collapse to a local minima more quickly.

### Convolutional Neural Network Simulations

We tested ACLS *in silico* on a deep convolutional neural networks with the following layer architecture:

Layer 1: Image input layer with 9 × 9 pixel size.

Layers 2-4: 3 feature extraction layers consisting of a) a convolutional layer with 8, 16, and 32 3-by-3 pixel filters for layers 2, 3, and 4, respectively (stride: L2:[1,1], L3-4:[2,2], with same padding), b) a batch normalization layer, c) a rectified linear layer, and d) a max pooling layer.

Layer 5: Fully connected layer.

Layer 6: Softmax classification layer.

Unless otherwise noted, the activity of the fully connected layer (e.g. the final hidden layer) was the response used to test ACLS.

CNNs were trained to classify numeric images using the MNIST dataset (7500 images of 0 to 9 digits; 750 samples from each digit; Figs. 2 and 3A-B). To decrease the dimensionality of the input, images were spatially downsampled to 9 × 9 pixels. The numeric information of the images was not lost with downsampling; the CNN’s performance was well above chance (92% on average). Unless otherwise noted, each block consisted of 5 repetitions of 10 different stimulation patterns.

To investigate the impact of network complexity on ACLS learning rate (Fig. 3E-F), a CNN was trained to classify 2,10,18 or 26 alphabets from EMNIST dataset (91650 images of 26 English alphabets; 3525 samples from each letter). In addition, a second fully connected layer was added prior to the classification layer to achieve similar classification accuracy (>87%) across all dimensions of inputs. ACLS was tasked with controlling the first fully connected layer. Similar to above, each block was consisted of 5 repetitions of 10 unique stimulation patterns. For these simulations, white Gaussian noise (µ = 0, σ = 0.5) was added to the response of the first fully connected layer.

### Surgical Procedures

Surgery was performed under isoflurane anesthesia (2.5% induction, 1.5% maintenance). Mice were given injections of dexamethasone (2 mg/kg), meloxicam (0.1 mg/kg), and buprenorphine (1 mg/kg) before surgery. A 3-4 mm diameter craniotomy centered at 2.2 mm lateral and 1 mm anterior to lambda was performed. An aluminum ring (Ziggy’s Tube and Wire) of the same diameter as the craniotomy was secured inside the window using glue (VetBond, 3M). A second smaller (0.5 mm) craniotomy was made contralateral to the first craniotomy, centered at 0.7 mm lateral of midline and 0.7 mm anterior to bregma. A custom-made ground wire secured to an Amphenol pin (A-M systems) was implanted into the second craniotomy and secured with glue (VetBond, 3M). A titanium head-plate for head restraint was also affixed to the skull of each mouse using dental cement (Metabond, Parkell). The first large craniotomy was kept moist with saline and surgical lubricant (Surgilube, HR Pharmaceuticals) and covered at the end of surgery and between recording sessions with silicone elastomer (Kwik-Sil, World Precision Instruments). Mice were given post-operative pain medication for the first 48 hours in 24-hour intervals and monitored for a 5 day recovery period

### Neural Recordings: Awake Acute

Acute recordings began at least 5 days following surgery and were carried out up to 4 weeks after surgery. Mice were habituated to the setup and handling 2-4 days before recordings began. For recordings, mice were head-fixed in a 1.5 inch diameter x 4 inch long polycarbonate tube.

For electrical stimulation and recording, a 64-channel silicon probe (Neuronexus A64 series; electrodes activated with iridium oxide to improve stimulation) was inserted in the primary visual cortex (Coordinates: ∼1mm anterior ∼2.2mm lateral of lambda) using a micromanipulator (Thorlabs Inc.). Extracellular signals were acquired using an RZ5 processor (Tucker-Davis Technologies) with a sampling rate of 25 kHz, filtered from 0.5 to 5 kHz. Electrical stimulation current was delivered using an IZ2 stimulator (Tucker-Davis Technologies) thorough 32 channels.

### Spike Counting and Detection

Action potentials (spikes) were detected by thresholding the filtered electrophysiological signal. Threshold levels were set as in Quiroga et al., 2004:

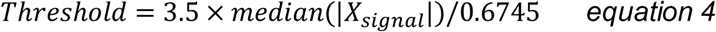

where *X*_*signal*_ was the filtered signal during the last 81ms of each trial in acute awake experiments. For adaptation experiments *X*_*signal*_ was the filtered signal during the last 5 seconds of the whole recording experiment. To allow for real-time, closed-loop control, we did not sort spikes into single neurons. As such, all spiking activity is multi-unit. To improve recording fidelity and decrease artifacts due to animal movement, one channel was selected as reference channel. The activity of this channel was subtracted from the rest of the recording channels.

The multi-unit response to electrical stimulation was quantified as the number of spikes during a 200ms window that excluded potential stimulation artifacts. This was done by setting the minimum possible starting point of spike count window to 10ms after the last stimulation pulse. This window was automatically adjusted for each experiment to the period with maximum information about the delivered electrical or visual stimuli. The mutual information *I(R; S)* between individual multi-unit response (R) and stimuli (S) was computed as in Tafazoli et al., 2017:

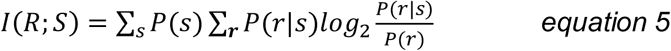

where *P(s)* was the probability of presentation of stimulus *s, P(r*|*s)* was the probability of observing response *r* upon presentation of stimulus *s*, and *P(r)* was the probability of observing response *r*. The onset of the spike counting window was the time point with the maximum mutual information, averaged across all stimulation channels.

### Selection of recording and stimulation channels

After determining the spike-counting window, the mutual information about stimuli (electrical or visual) was calculated as in equation 5. Multi-units that conveyed more than 0.05 bits of information about the identity of the stimulus were initially selected as ‘responsive’. Activity of these units was visually inspected by the experimenter and those that did not have waveform characteristics of a neuron were omitted. Stimulation channels were then selected as the set of channels with the closest distance to target sites.

### Visual stimuli

For *in vivo* experiments, visual stimuli were taken from the Caltech 101 database (Fei-Fei et al., 2004). Between 4 and 20 images were used to generate the response space (the 10 images used for the Fig. 4 example are shown in Fig. S5A). The stimuli were displayed on a 23-inch LCD monitor (LG Flatron234V), with 1920 x 1080 pixel resolution, 60 Hz refresh rate, 14ms response time, 250 cd/m2 maximum brightness). The monitor was tilted 20 deg on the elevation axis and 60 deg on the azimuth axis and was positioned 10 cm from the animal’s right eye. Thus, the monitor spanned a visual field of 137 deg azimuth and 70 deg elevation. To map the visual response space, visual stimuli were randomly presented in a sequence (200ms stimulus on, followed by 1000ms of blank gray screen). The intensity of the gray screen matched the average luminance of the presented images.

### Adaptation experiment

Adaptation experiments immediately followed a subset (n=4) of the awake recording experiments. Electrode placement, electrical stimulation, and neural recordings were all performed as in awake acute experiments.

Prior to adaptation experiments ACLS was used to learn two electrical stimulation patterns (ES1, ES2) that reproduced the population-level response of two target responses of visual stimuli (VS1 and VS2; learning was performed as in awake experiments). ES1 and ES2 were taken as the stimulation patterns during the last block of learning with the smallest error to the target response of visual stimuli VS1 and VS2, respectively.

Each trial of the adaptation paradigm consisted of two visual stimulus presentations (either VS1 or VS2), followed by electrical stimulation (either ES1 or ES2). This yielded eight different trial types: VS1→VS1→ES1, VS2→VS2→ES1, VS1→VS2→ES1, VS2→VS1→ES1, VS2→VS2→ES2, VS1→VS1→ES2, VS1→VS2→ES2, VS2→VS1→ES2. All stimuli were followed by a 100ms interstimulus interval (ISI). Trials were separated by 4500-5000ms intertrial interval (ITI). ISI and ITI values were chosen based on previous studies (Jin et al., 2019). Each trial type was repeated 30 times and the trial order was random.

## Quantification and Statistical Analysis

### Statistical Analysis

All analyses were performed in MATLAB (2017a-2019b, Mathworks). Number of mice used was based on previous work (Zhang et al., 2018). All statistical tests and associated values are reported in the main text.

### Drift Correction

To quantify global drift in the response space over time in the *in vivo* experiments, the initial stimulation pattern of each run was repeated twice per block. Drift was measured as the Euclidean distance between these repetitions and the target response (Fig. 4G-H, purple and orange dotted lines). Because these stimuli were sampled infrequently, the moving mean of this drift was computed across 6 blocks. The error of these repetitions was not reported to the ACLS algorithm. To correct for this drift, the error of ACLS evoked patterns was divided by the drift for each block (Fig. 4I-J).

### Quantifying Noise

To quantify noise in the response to electrical stimulation, each stimulation pattern was repeated 5 times per block during *in vivo* experiments. Noise was computed as the pairwise Euclidean distance between repetitions of the same stimulus, averaged across all stimuli per block.

### Quantifying ACLS Success

Success of each ACLS recording was quantified in two ways. First, a decrease in error from the first to the last block was computed as: Δ*ε = ε*_*start*_ − *ε*_*end*_ where *ε* is the error function (*equation 1*) during the first and last block. Δ*ε >* 0 was considered a success.

Second, a decrease in error over time was determined by fitting the first order exponential, *f(x) = ae*^*λx*^ + *c* to the error trajectory. A negative value of λ (e.g. a decreasing function) was considered a success.

### *Q*uantifying ACLS Selectivity

We sought to test the ability of ACLS to selectively evoke a population-level neural response. Selectivity was calculated as *s = ε*_*run*1, *target*1_ − *ε*_*run2, target1*_, where *ε*_*run*1, *target*1_ is the error of the current ACLS run to its target, and *ε*_*run2, target1*_ is the error of the other, simultaneous (and independent), ACLS run to the same target. Alternatively, we calculated selectivity as *s = ε*_*run*1, *target*1_ − *ε*_*run1, target2*_ which tested whether a run was closer to its target than the other target (error was drift-corrected to compensate for a global shift towards either target). In both cases *s* < 0 was considered a success. These errors captured specificity of ACLS and ensured the reduction in error was not due to a general collapse to a common response space across runs.

### Logistic Regression

Stepwise logistic regression was used to quantify the impact of *in vivo* experimental features on ACLS success and selectivity rates. AIC was used as the inclusion criteria for model terms. For each success or selectivity metric, stepwise models were fit using the following predictors: signal to noise, magnitude of shift, travelable distance, and average firing rate in response to all stimulation in the last block of learning (details below). These predictors were chosen because they capture a diversity of possible influences on ACLS success.

Signal to noise ratio was calculated as 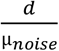 where *d* is the Euclidean distance (in 2D space; see *equation* between the two selected targets in a recording, and *µ*_*noise*_ is the average noise during the run. Magnitude of shift was calculated as 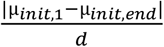 where *µ*_*rep*_ is the mean distance between responses to repetitions of the initial stimulus pattern and the target response (e.g. purple and orange dotted lines in Fig. 4G). Subscripts *1* and *end* denote first and last block, respectively. *d* is the distance between targets as above.

Travelable distance was calculated as 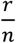 where *r* is the average distance between evoked and target responses and *n* is the noise, both taken during the first block of learning.

Average firing rate was calculated as the mean firing rate in response to all stimuli (including repetitions of the initial stimulus patterns) during the final block.

For all predictors, distance was computed in the 2D PC space. As described in the main text, this analysis revealed that signal to noise ratio was predictive of selectivity (p=0.018). Regressions for the remaining metrics failed to reject the null hypothesis (p of final model >0.1)

### Adaptation Index

The adaptation effect was quantified using an adaptation index, 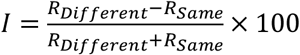 where *R*samewas the average multi-unit activity in response to a repeat of the same stimulus and *R*_*Different*_ was the average multi-unit activity in response to a different stimulus. Activity was averaged in 50ms windows: 0ms-50ms, 50ms-100ms, 100ms-150ms, 150ms-200ms and 200ms-250ms, all relative to the onset of stimulation.

## Supplementary Information

### Supplemental Methods: *in vivo* anesthetized mouse recordings

#### Supplemental Results: ACLS control of different CNN hidden layers (Fig. S1)

We tested the ability of ACLS to control representations at different depths of a CNN. Here, we controlled representations in the 1^st^ convolutional layer, 2^nd^ convolutional layer, and the fully connected layer of the CNN (Fig. S1a; CNN architecture was the same as described in the main text, see Methods for details). Each layer was controlled in a separate run, but the same instance of the CNN was used for all runs. We began each experiment by mapping the response of the target layer (e.g. convolutional layer 1) to 1000 images of numbers 0, 1, 2, and 3 (250 images of each number). A ‘target’ response was randomly selected from the cluster of responses to images of 1. ACLS was then tasked with learning to produce this target response in either the reduced, two-dimensional, PC space or in full-dimensional space. As seen in Figure S1, the ACLS was able to minimize the error at each layer, in both the full-dimensional space (Fig. 1B-D) and in the reduced-dimensional space (Fig. S1E-G).

Note, that the learned stimulation patterns closely match the original target when learning to reproduce responses at early layers (Fig. S1B and C). However, at deeper layers, ACLS found adversarial images that were different from the original stimulus input, but still generated the same response in the target layer (Fig. S1D). This shows that the adversarial images are not a by-product of the ACLS approach. Instead, the adversarial images are a consequence of the highly convergent nature of CNNs (Szegedy et al., 2014).

In addition, it is important to note that the efficacy of learning in a reduced-dimensional space depends on how much of the variance in activity is captured by that space. In early layers, the dimensionality of the representation was high and so the first two PCs captured a limited amount of the variance (<30%, Fig. S1E, right bar plot). In contrast, the last layer had a lower dimensional representation (∼70% of variance explained by the first two PCs, Fig. S1G, right bar plot). Although ACLS was able to learn in the reduced-dimensional space for both layers (Fig. S1E-G), this learning transferred to the full-dimensional space better in the deeper layers (because of the lower dimensionality of these representations, Fig. S1E-G). This is also reflected in the fact that the stimuli generated by the ACLS algorithm learning in the reduced-dimensional space were different from the stimuli learned in the full-dimensional space for early layers (e.g. compare S1B to S1E) but not deeper layers (e.g. compare S1D to S1G).

#### Supplemental Results: *in vivo* anesthetized mouse recordings (Figures S3-S4)

Supplemental experiments tested the ability of ACLS to control neural populations in the primary visual cortex of anesthetized mice. These experiments closely followed the methods of *in vivo* awake mouse recordings.

5 male and 5 female mice were used for these experiments. Acute surgeries closely followed the main text with the exception that craniotomies were 2mm in diameter and no metal ring was implanted. Post-surgery, mice were transferred to the recording rig and maintained under isoflurane (1.5%) or ketamine/xylazine (87.5mg/kg Ketamine and 12.5mg Xylazine every 30 minutes, continuous intraperitoneal infusion) anesthesia. As in awake acute recordings, a 64-channel electrode was lowered into visual cortex to allow for stimulation and recording. Selection of recording and stimulation channels closely followed awake recordings: in brief, at the beginning of each recording a predefined set of charges (3, 5, and 8nC) was applied to each stimulation site (separately) while recording activity across all sites. Responsive multi-units were detected by the same methods used for awake recordings. To find the effect of each channel on each multi-unit, a linear regression model was fitted according to equation S1:

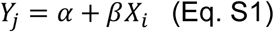

where *X*_*i*_ were stimulation values for channel *i* and *Y*_*j*_ were response from multi-unit *j. β* values were clustered with a hierarchical clustering algorithm to find the minimum set of stimulation channels that could effectively stimulate all target multi-units. This set of stimulation channels was used for the subsequent ACLS learning experiment. The ACLS algorithm was not provided with these stimulation patterns, responses, or model fits.

The objective of these supplemental experiments was to test the ability of ACLS to learn to produce arbitrary electrically-evoked neural population responses (Fig. S3). To this end, we mapped the response space to 10 random electrical stimulation patterns (Fig. S3A-B). Each pattern was repeated 20-30 times; and the response of the network to each pattern repetition was taken as the number of spikes in a 200 ms window, starting after the stimulation artifact and chosen to maximize information about the stimuli (see Methods for details). As in awake experiments, we projected these responses onto the first two principal components of the entire distribution (see Fig. S3C gray dots for example).

As in awake experiments, the response pattern of two of these patterns were selected manually. This was done blind to the actual stimulation pattern but in a way that the responses were separated in the 2D space. Two simultaneous ACLS runs were then tasked with crossing the response space to move between these two targets (Fig. S3C). Each run was initialized with the stimulation pattern that evoked one target and learned to reproduce the response of the *other* target (e.g. orange to purple and vice versa in Fig. S3C). Figure S3D-E shows that ACLS decreased the error of evoked responses for both runs in the reduced and full dimensional space. Drift correction and error calculations were performed as for awake animals. However, it is important to note that the constraint that ACLS runs must switch positions mitigates the impact of systematic drift in responses.

Figure S4A shows the error trajectory of ACLS across 68 runs during 34 recordings in 10 anesthetized animals. ACLS successfully reduced the error of the evoked response in 76.47% of runs (52/68 runs, p = 10^−6^, Binomial test). On average, the error of evoked responses decreased by 30.79% in block 10 and by 49.32% in block 15, approaching the noise floor over time (Fig. S4B). The evoked responses were selective for the target response pattern on 79.41% of runs, as measured by testing if the error of the current run to its target was less than the error of the other simultaneous run to the same target (54/68 runs, p = 10^−7^, Binomial test). Success and selectivity were calculated as in awake animals (see Methods). As noted above, because each target was initialized at the other target location, learning could not be explained by drift in the system.

#### Supplemental Results: Additional example ACLS runs in awake mice (Fig. S7)

Figure S7 shows additional examples of ACLS learning electrical stimulation patterns that reproduce visually-evoked target activity. All experiments follow the procedures outlined in the main text and methods and used for Figures 4-5.

These examples highlight some of the diversity of ACLS runs observed *in vivo.* Fig. S7A-B demonstrates a recording where both ACLS runs started close to one of the target responses (purple star). Here, one of the runs is tasked with staying close to this target (which it does; purple line) while the other run (orange line) traverses the low-dimensional response space toward its respective target (orange star). A similar trajectory pattern is observed in the experiment shown in Fig. S7C-D.

Conversely, Figure S7E-F shows an example recording in which the two runs start closer to the others’ target response, and then traverse the response space towards their target response. In addition, this example highlights how one of the runs (orange) starts near its noise floor (dotted line in F) and thus is limited in its ability to appreciably decrease its error.

Finally, example Figure S7G-H shows two simultaneous runs with initial, electrically-evoked responses that reside well outside the visual response space (gray dots). Over time, the learned stimulation patterns elicit responses within the visual response space and selectively approach each target (similar to Fig. 4).

#### Supplemental Results: Quantifying drift in an anesthetized mouse (Fig. S8

Supplemental experiments were performed to characterize the drift in neural responses over an extended time period. Experimental methods followed those of the anesthetized ACLS experiments (detailed above). However, instead of using the ACLS system, we repeatedly delivered a fixed set of 10 electrical stimulation patterns over ∼82 minutes (240 repetitions of each pattern). Stimulation patterns were delivered over all 32 stimulation electrodes. Patterns were delivered in random order. On average, a stimulation was delivered approximately every 2.1 seconds. To measure the drift in the evoked response, only the response of stimulation-responsive multi-units was used in all subsequent analyses (see Methods for details). The response of this population of multi-units was then projected into a low-dimensional space defined by the first two principal components.

Figure S8A shows the trajectory of the population response to three example stimulation patterns, smoothed over time (red, green and blue trajectories; color saturation increases with time). Importantly, these three patterns reside in different parts of the low-dimensional space, reflecting the fact that the neural population responded differently to different stimulation patterns (consistent with our ACLS experiments). However, it is also clear that there is significant drift in the response to the same stimulation pattern over time. This drift was seen in the response to all 10 stimulation patterns (Fig. S8B).

To quantify drift, we measured the variance in the response to the same stimulation pattern (calculated as the Euclidean distance in the low-dimensional space). This was done in 50 trial windows, allowing us to quantify variance in the response over time. As seen in Figure S8C, the variability in the response changed over time. Interestingly, the variance in response was correlated across the different stimulation patterns (average rho=0.57; averaged across pairwise comparisons between the 10 stimulation patterns; p<0.05 in 43 of the 45 possible pairwise comparisons between stimulation patterns). This suggests drift was due to system-wide changes. Under anesthesia, these state changes seemed to happen regularly; the autocorrelation of the variance in response had a semi-periodic structure suggesting states fluctuated every ∼10-15 minutes (Fig. S8D).

###### Box S1.

**Related to Figure 1. Pseudocode of ACLS framework**

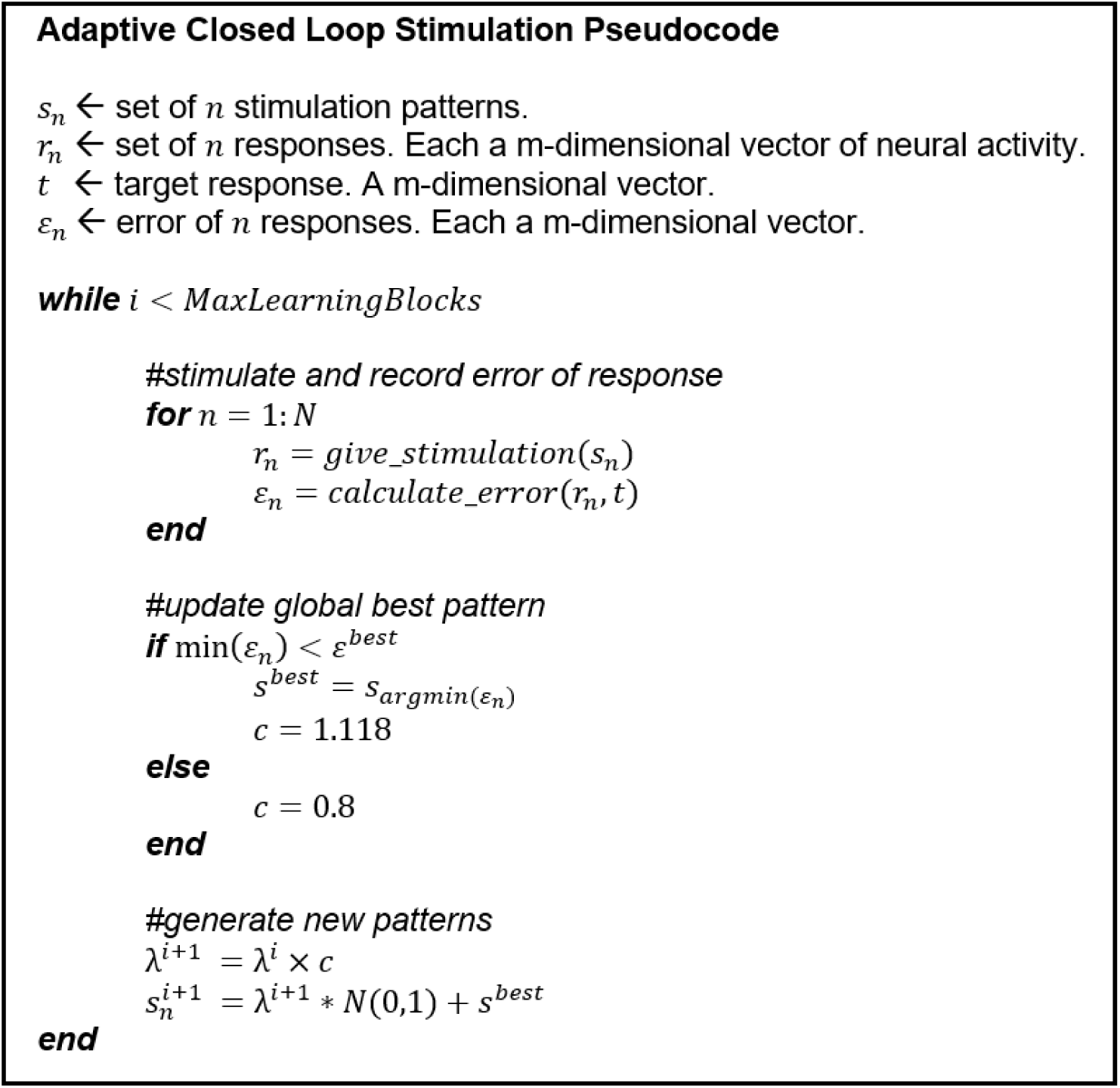

**Figure S1.**
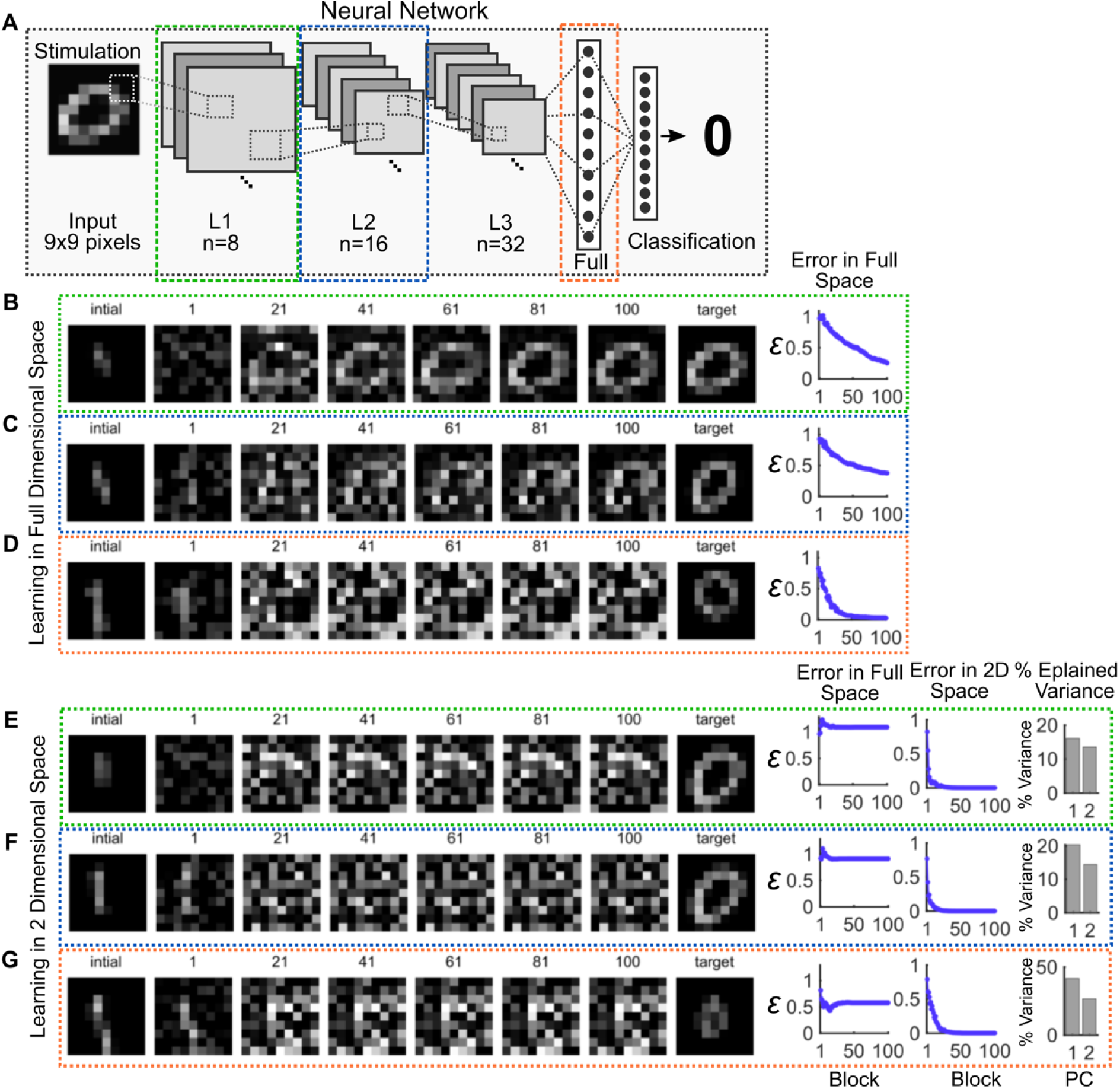
Related to Figure 2. ACLS-learned stimulation patterns when controlling different CNN layers. **(A)** ACLS was tasked with controlling different hidden layers of a CNN (colored boxes) by reducing error in either the full dimensional or reduced dimensional space. Display follows Figure 2A. **(B-G)** Example evolution of learned stimulation patterns when calculating error in **(B-D)** full or **(E-G)** reduced dimensional space. Errors were normalized to distance between initial point and target points in full or reduced dimensional space. Numbered images show the best learned stimulation pattern during a subset of 100 blocks (each block consisted of 1 repetition of 20 unique stimulation patterns). For these simulations, no noise was added to responses of fully connected layer. “Initial” shows the initial stimulation pattern provided to ACLS. “Target” shows the pattern originally used to generate the target response in the network. As seen in B-C, the patterned learned by ACLS was similar to the original pattern when controlling the earlier layers. For display purposes, the image intensity was normalized to the minimum and maximum for each example run (i.e. within a row).

**Figure S2.**
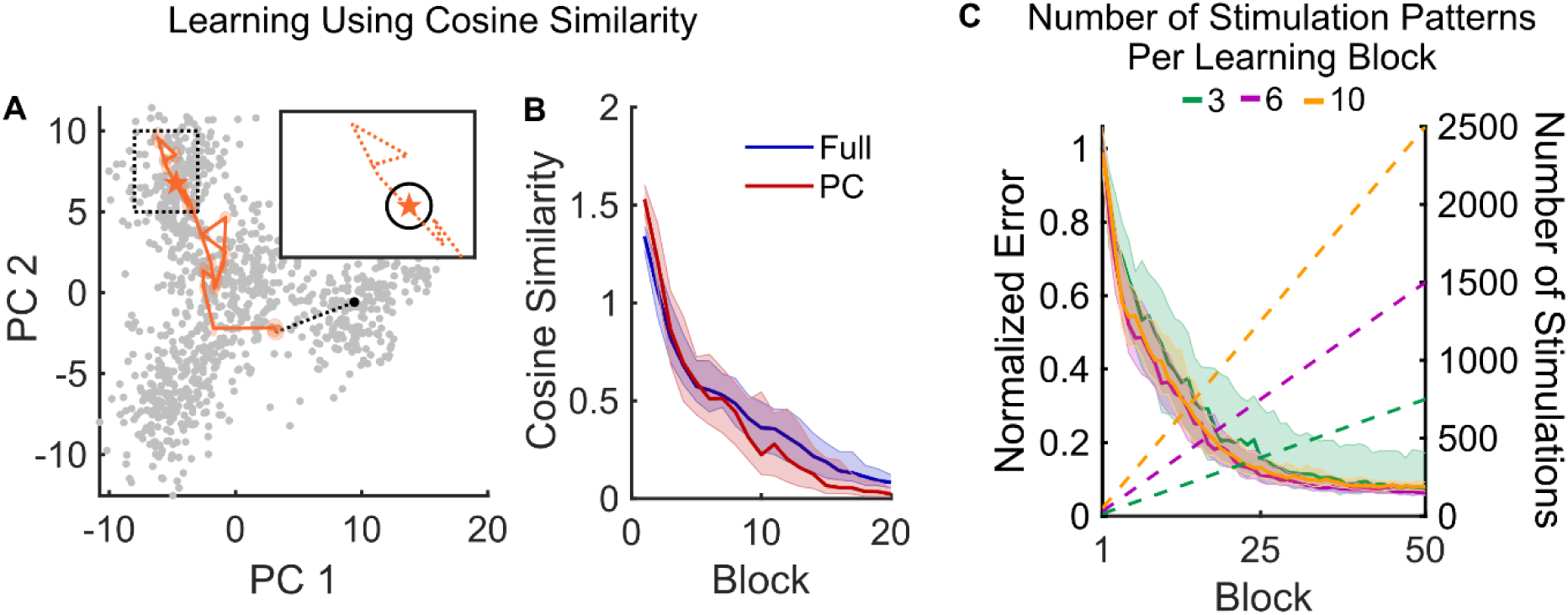
Related to Figure 3. Testing additional ACLS parameters. **(A)** Example trajectory of ACLS learning to control the final hidden layer of CNN using a cosine similarity error function. The error function was *ε =* 1 − *cos θ* where 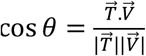 and *T* and *V* were the target and current response vectors, respectively. Display follows Figure 2F. **(B)** ACLS decreased the cosine similarity error function in both full (blue) and two-dimensional PC space (red). Lines and shaded regions show the median and IQR of 50 replications. **(C)** Varying the number of stimulation patterns per block impacted ACLS learning. Learning was quantified when using 3 (green), 6 (purple), or 10 (orange) unique stimulation parameters per block. Solid lines show median error of evoked responses over learning with shaded regions showing IQR over N=50 replications. Higher number of stimulation patterns per block resulted in more reliable learning (demonstrated by a lower variance in the learning curve), but required more total stimulations, shown as an increase in the cumulative number of trials across blocks (dashed lines, right y-axis).

**Figure S3.**
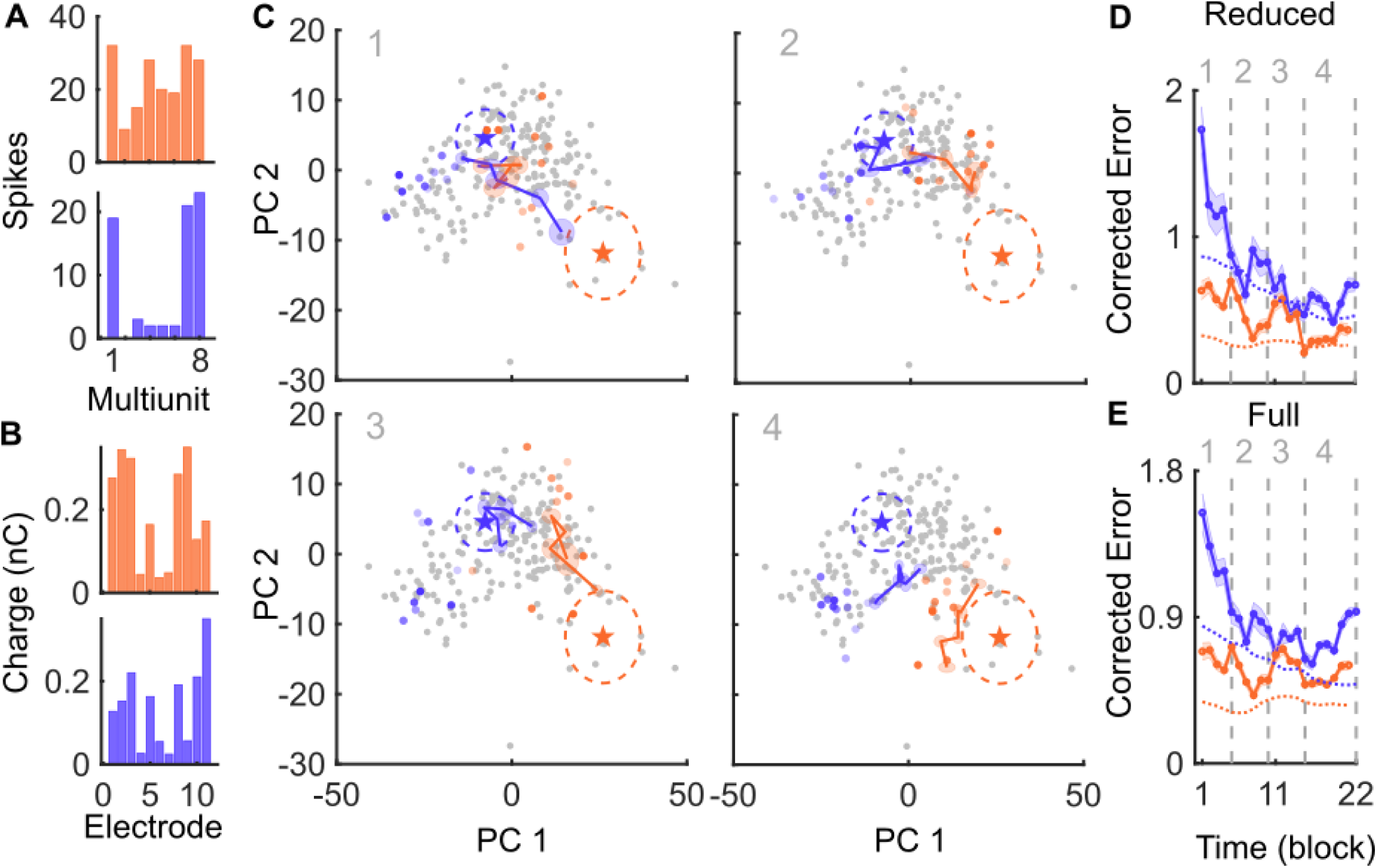
Related to Figures 4-5. ACLS learns to produce specific patterns of neural activity in vivo anesthetized mouse primary visual cortex. **(A)** Example multi-unit activity in response to random electrical stimulation in the primary visual cortex of an anesthetized mouse (see Supplemental Methods for details). Display follows Figure 4B. **(B)** The electrical stimulation patterns that evoked the corresponding colored responses in A. **(C)** Example trajectory of ACLS learning to reproduce arbitrarily-defined population responses (previously elicited by random electrical stimulation). Gray dots show the responses to random electrical stimulation projected into a 2D space defined by the first two principal components (as in Figure 4C). Two responses were selected as target responses for ACLS to achieve (orange and purple stars; corresponding to firing and stimulation patterns shown in A-B). ACLS was tasked with learning to reproduce these responses, starting from the stimulation pattern that produced the response of the opposite target (e.g. purple run starts at orange and learns towards purple, and vice versa). Panels in C show the learning trajectory of the algorithm across 22 blocks. Display follows 4C. **(D)** Corrected error over time. Display follows Figure 4G and 4I.

**Figure S4.**
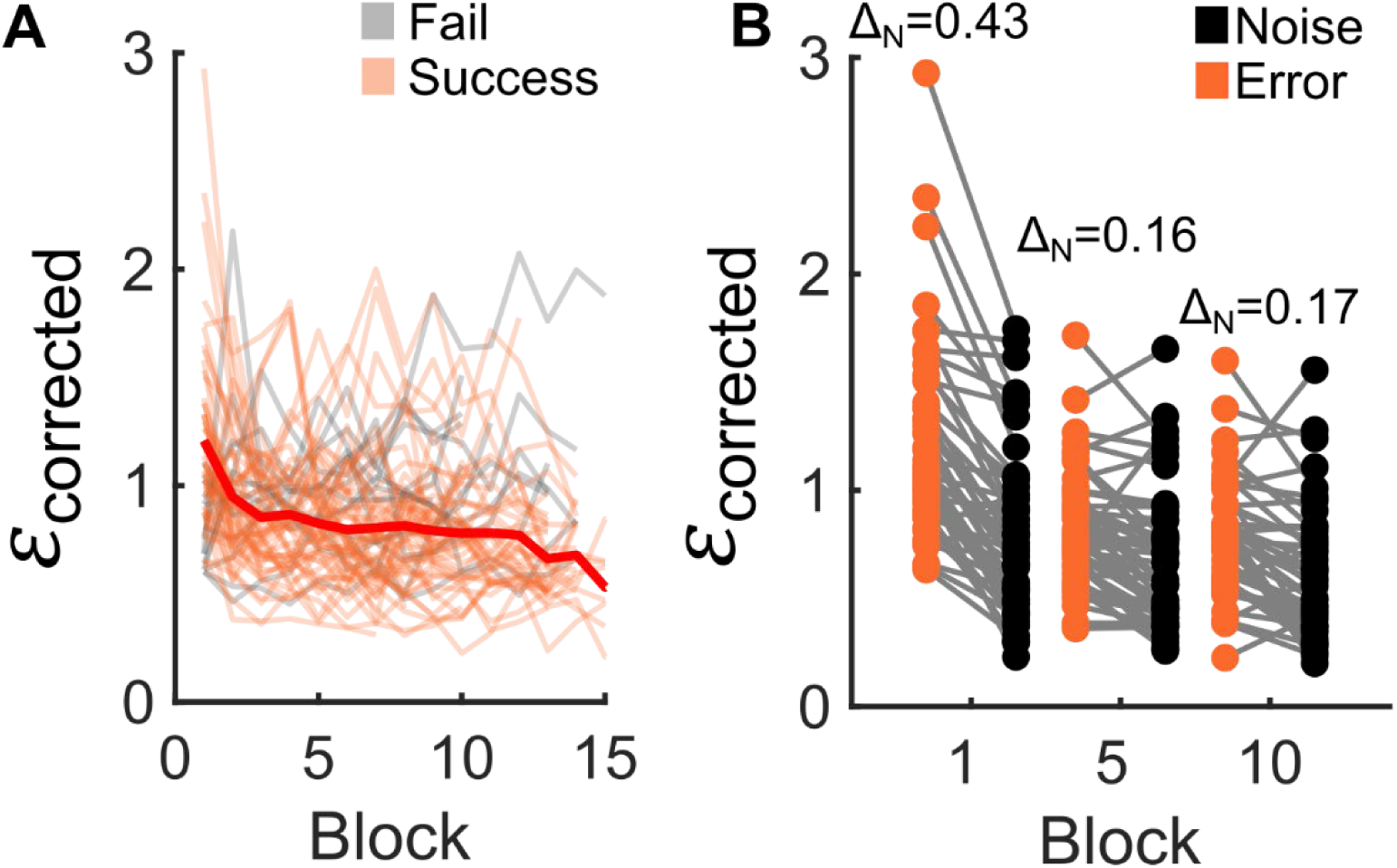
Related to Figures 4-5. Quantifying ACLS success across all anesthetized recording sessions. **(A)** Corrected error over time across N=34 runs in N=68 recordings in N=10 anesthetized mice. Display follows 5A. Figure cropped to 15 blocks to allow for visualization of all ACLS runs. N=5 continued to decrease their error after this point. **(B)** Pairwise comparisons of error and noise levels at three time points. Display follows Figure 5B.

**Figure S5.**
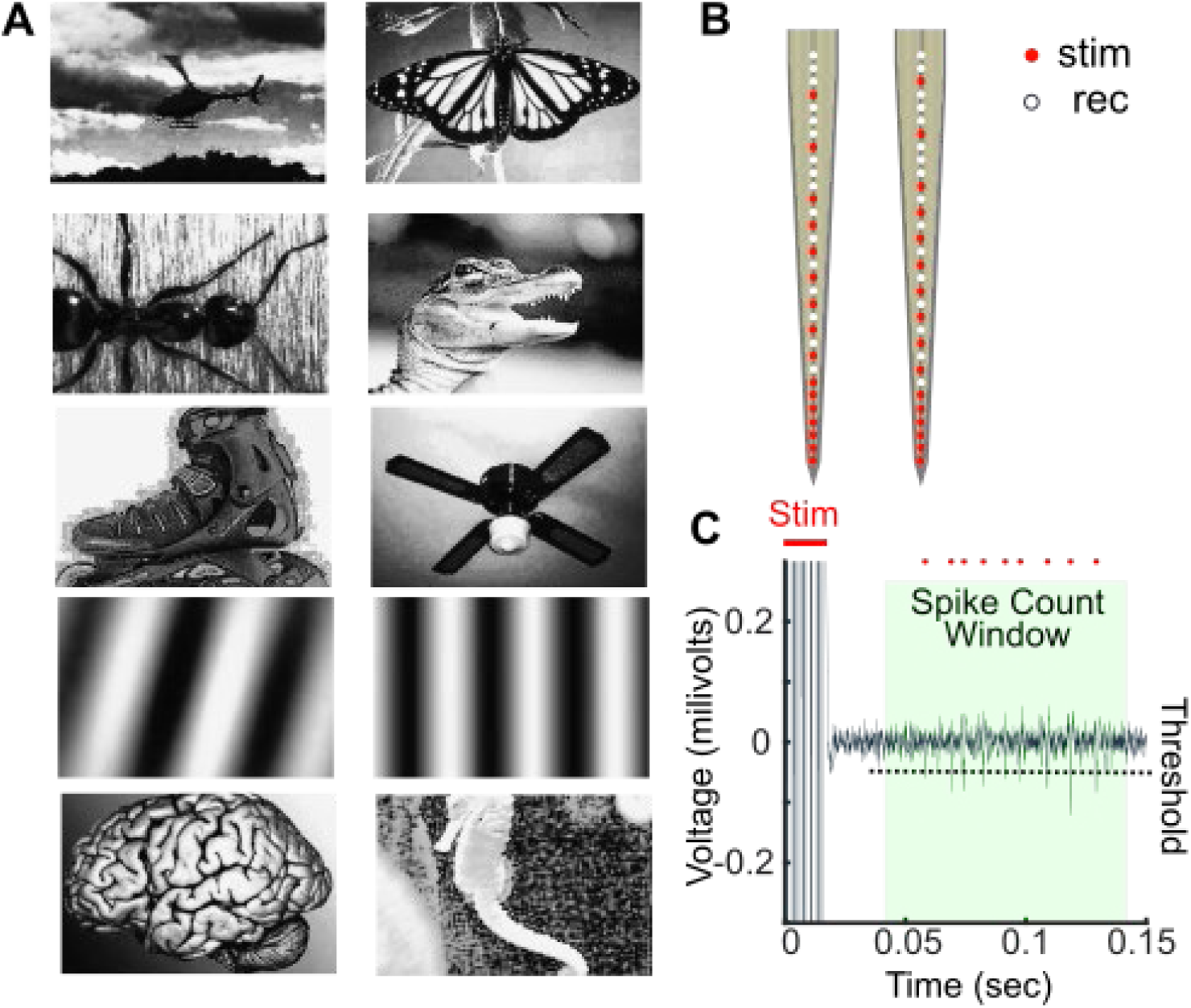
Related to Figures 4-5. Additional Details Related to Methods. **(A)** Visual stimuli from Caltech 101 database used in the example recording shown in Figure 4. **(B)** Schematic of 64 channel silicone probe used in awake recordings. Red and white sites denote stimulation and recording electrodes, respectively. **(C)** Schematic of multi-unit spike detection. Threshold detection (black dotted line) was used to detect multi-unit spikes (red dots) during a predefined window (green box). This window excluded stimulus artifacts (red line).

**Figure S6.**
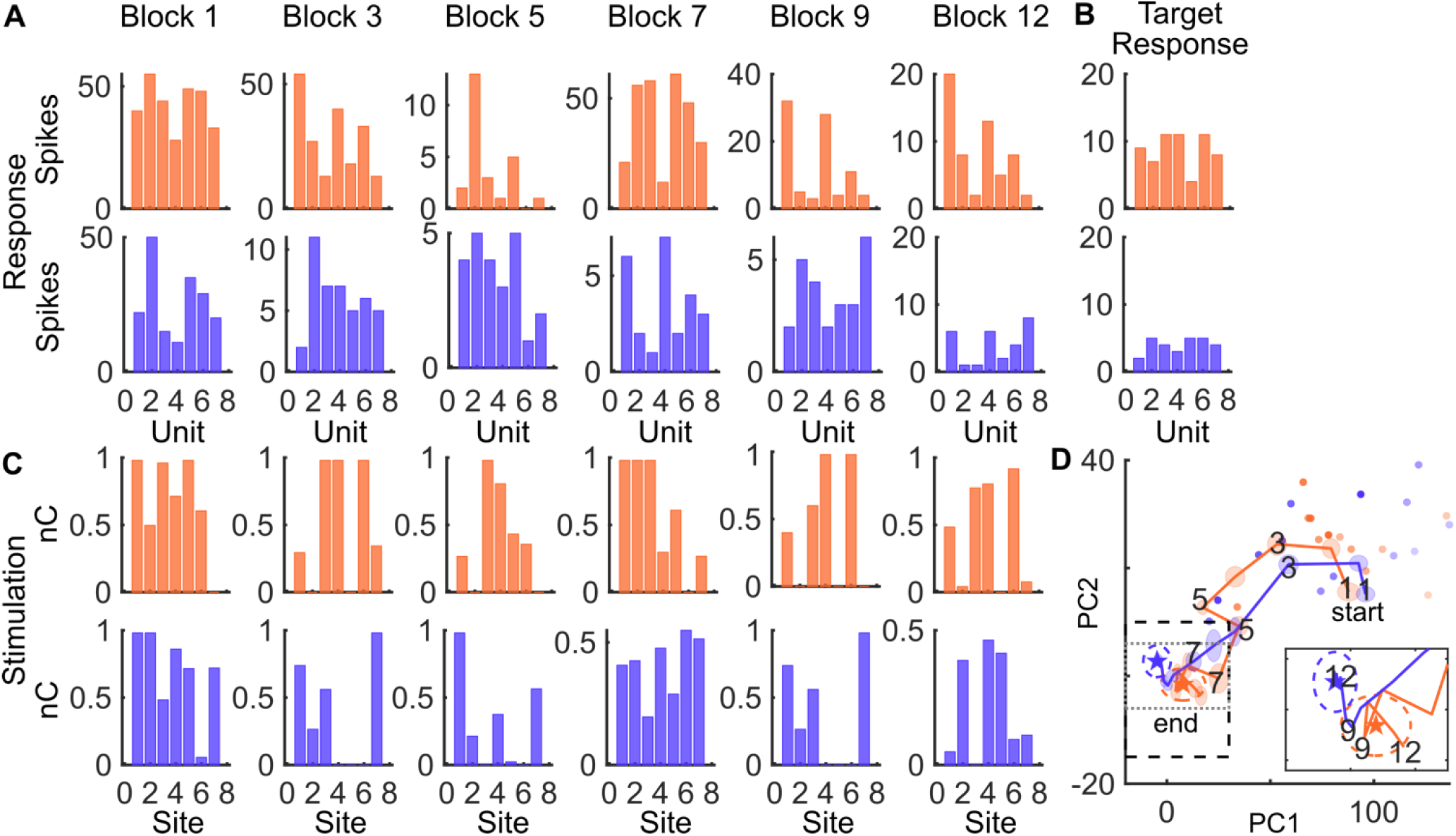
Related to Figure 4. Example stimulations and responses during ACLS learning. **(A)** Multi-unit responses during ACLS learning for the example session shown in Figure 4. Each plot shows the best response during a given block (e.g. the response with the lowest error). Orange and purple denote responses for the two simultaneous ACLS runs. Note: the y-axes are independently scaled in order to show the full dynamic range of response values. **(B)** Target responses for each run. Colors match panels A and C **(C)** Stimulation pattern that evoked the corresponding responses in A (aligned by column). **(D)** Learning trajectory in PC space associated with A-C. Numbers correspond to example blocks in A and C.

**Figure S7.**
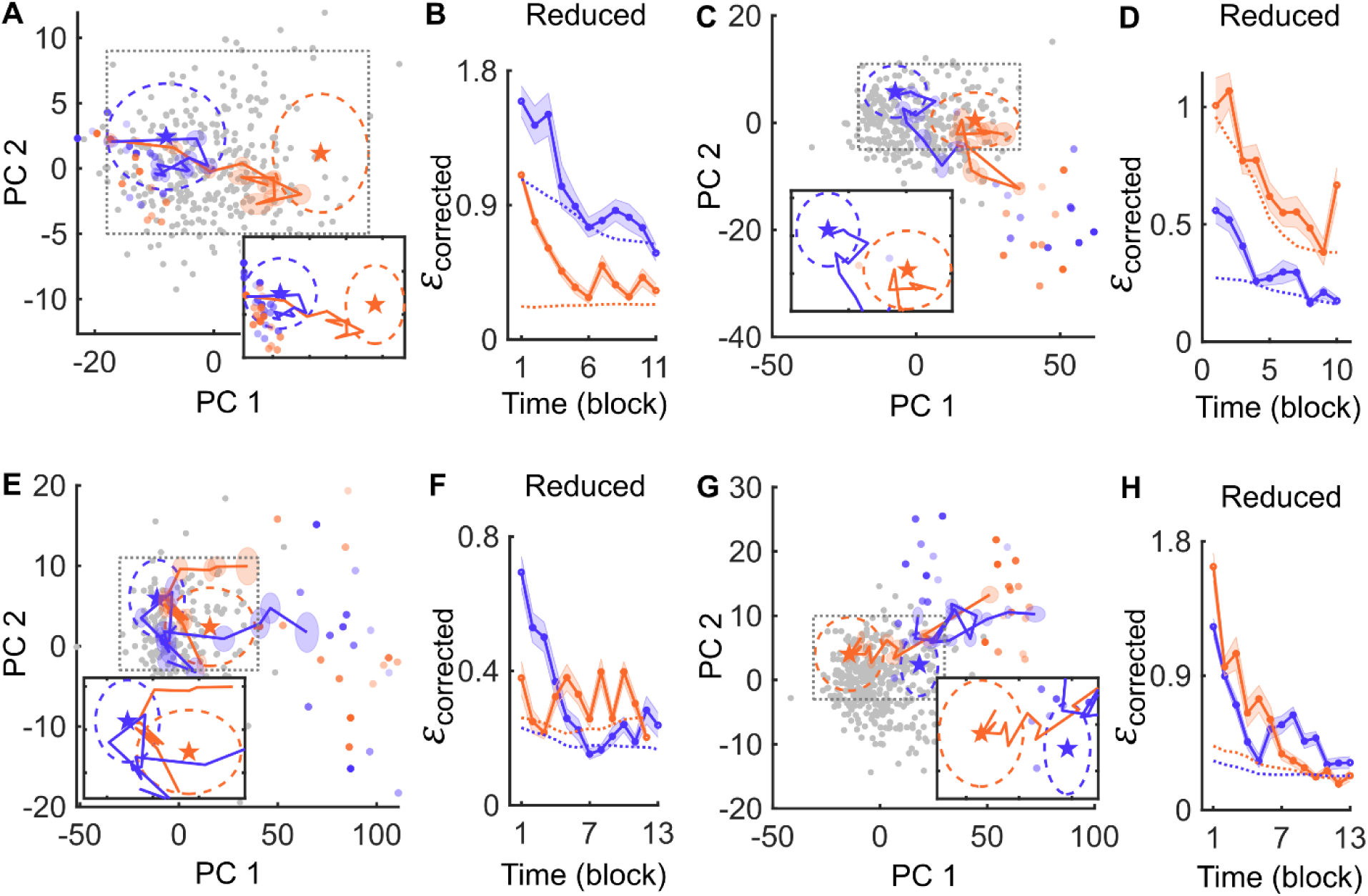
Related to Figures 4-5. Additional examples of ACLS runs. **(A-G)** The left panel of each pair of panels shows the **(A, C, E, G)** 2D trajectory of example ACLS runs in awake mice and the **(B, D, F, H)** associated drift-corrected error across blocks. The panels follow the displays in Figure 4G and 4I for the left and right panels, respectively.

**Figure S8.**
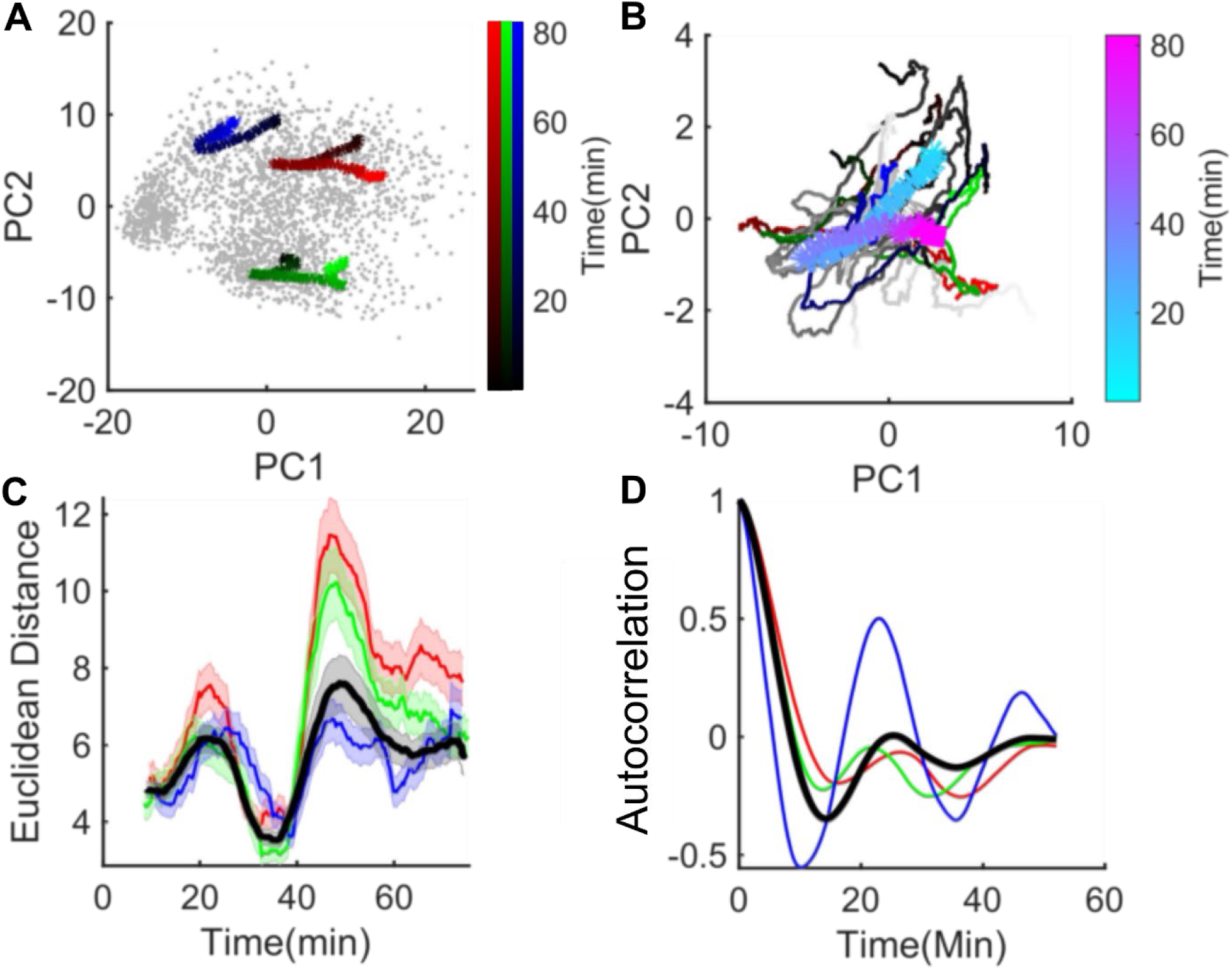
Related to Figures 4-6. Quantifying drift *in vivo* over long periods of time. **(A)** Low dimensional projection of activity of 10 multi-units recorded while 10 random stimulation patterns (32 channels) were delivered 240 times each over 82 mins (gray points). The mean trajectory over time for three example patterns is shown in red, blue and green. Color gradient shows progression over time, with more saturated timepoints being later in the session. **(B)** Mean centered trajectories of responses to all 10 patterns. Red, blue and green show example patterns from A. The remaining patterns are shown in gray. Average of all trajectories is shown in the thick line, running from cyan to purple over time. **(C)** Time-averaged pairwise Euclidean distance between responses in the 2D response space. Red, blue, and green trajectories correspond to the 3 example patterns in A. Black curve shows the mean values across all 10 patterns. Shaded region shows SEM of N=50 averaged time points. **(D)** Autocorrelation of pairwise Euclidean distance computed in C. Red, green, and blue lines show example trajectories. Black line shows the mean across all 10 trajectories.

**Table S1.** ACLS parameters and experimental results for individual awake experiments. Each recording (row) included 2 runs. Run-specific values are denoted in column labels. See methods for descriptions of parameters. Decrease in error was calculated as: Δ*ε = ε*_*start*_ − *ε*_*end*_. Thus, positive values denote ‘successful’ runs. Selectivity was calculated as *s = ε*_*run*1, *target*1_ − *ε*_*run*2, *target*1_.Thus, negative values denote run selectivity for its target stimulus.

| Recording # (2 runs per recording) | # Multi-Units | # Stimulation Sites | # Unique Stimulation Patterns per Block | # Repetitions of Each Stimulation Pattern | SNR (PC Space) | # Blocks (Run 1) | Blocks (Run 2) | Decrease in Error: Run 1 (Corrected PC Error) | Decrease in Error: Run 2 (Corrected PC Error) | Selectivity: Run 1 (Corrected PC Error) | Selectivity: Run 2 (Corrected PC Error) | Selectivity: Run 1 (Corrected Real Error) | Selectivity: Run 2 (Corrected Real Error) |
| --- | --- | --- | --- | --- | --- | --- | --- | --- | --- | --- | --- | --- | --- |
| 1 | 5 | 10 | 6 | 5 | 0.71 | 14 | 14 | 0.36 | 0.26 | -0.62 | -0.13 | -0.29 | -0.03 |
| 2 | 5 | 4 | 6 | 5 | 0.59 | 13 | 14 | 0.22 | 0.98 | 2.15 | -1.53 | 1.69 | -1.93 |
| 3 | 11 | 10 | 6 | 5 | 0.96 | 13 | 13 | 0.50 | 0.96 | 2.50 | 5.32 | -0.41 | 5.02 |
| 4 | 6 | 16 | 6 | 5 | 2.53 | 13 | 13 | 1.75 | 0.35 | -0.15 | -0.76 | -0.67 | -0.03 |
| 5 | 7 | 6 | 6 | 5 | 1.85 | 13 | 13 | 0.53 | 0.16 | -13.70 | -2.74 | -14.24 | 1.55 |
| 6 | 8 | 6 | 6 | 5 | 1.18 | 13 | 13 | 0.29 | -0.71 | -0.83 | -1.98 | -0.35 | -1.44 |
| 7 | 6 | 7 | 6 | 5 | 2.35 | 14 | 15 | 0.04 | 0.69 | -9.21 | -26.91 | -7.90 | -27.39 |
| 8 | 6 | 6 | 6 | 5 | 2.93 | 13 | 13 | 0.46 | 0.18 | -20.41 | -12.04 | -17.18 | -11.41 |
| 9 | 7 | 7 | 6 | 5 | 1.07 | 12 | 12 | 0.71 | 0.41 | -19.69 | 2.54 | -22.68 | 8.73 |
| 10 | 7 | 10 | 6 | 5 | 1.53 | 13 | 12 | 0.46 | 0.18 | -3.72 | -15.80 | -5.05 | -14.69 |
| 11 | 5 | 5 | 6 | 5 | 1.45 | 13 | 13 | 0.76 | 0.32 | -4.24 | -4.15 | -3.63 | -3.90 |
| 12 | 5 | 6 | 6 | 5 | 1.02 | 12 | 12 | 1.19 | 0.58 | 12.42 | -15.57 | 4.91 | -12.21 |
| 13 | 10 | 9 | 6 | 5 | 1.28 | 14 | 15 | -0.53 | 0.23 | 12.87 | -28.30 | 12.20 | -27.03 |
| 14 | 6 | 8 | 6 | 5 | 2.19 | 10 | 10 | 0.38 | 0.34 | -29.95 | -5.80 | -29.99 | -0.73 |
| 15 | 8 | 10 | 6 | 5 | 2.98 | 13 | 13 | 0.91 | 1.40 | -14.39 | -24.21 | -9.76 | -22.40 |
| 16 | 5 | 7 | 6 | 5 | 3.99 | 11 | 11 | 1.02 | 0.77 | -13.98 | -12.35 | -13.32 | -8.29 |
| 17 | 7 | 16 | 6 | 5 | 2.48 | 16 | 16 | 0.17 | 0.38 | -6.02 | -11.79 | -5.60 | -8.62 |
| 18 | 5 | 5 | 6 | 5 | 1.10 | 12 | 13 | 0.67 | -0.08 | -7.47 | 2.21 | -3.24 | 2.38 |
| 19 | 7 | 7 | 6 | 5 | 0.76 | 11 | 10 | 0.31 | 0.96 | 9.48 | -12.49 | 5.52 | -7.68 |
| 20 | 5 | 13 | 6 | 5 | 0.86 | 14 | 14 | -0.53 | 0.00 | 14.79 | 2.79 | 5.87 | 9.71 |
| 21 | 8 | 9 | 6 | 5 | 0.45 | 9 | 9 | 0.28 | 0.00 | -9.34 | 12.78 | -8.68 | 13.22 |
| 22 | 10 | 16 | 6 | 5 | 1.23 | 19 | 20 | 0.47 | 0.59 | 2.81 | -3.96 | 2.15 | -1.67 |
| 23 | 7 | 16 | 6 | 5 | 2.89 | 10 | 10 | 0.01 | -0.01 | 1.73 | -2.47 | -1.64 | -1.98 |
| 24 | 9 | 10 | 6 | 5 | 1.30 | 12 | 12 | 0.45 | 0.03 | -15.07 | -0.63 | -17.00 | 3.02 |

